# Comparison of fine-scale recombination maps in fungal plant pathogens reveals dynamic recombination landscapes and intragenic hotspots

**DOI:** 10.1101/158907

**Authors:** Eva H. Stukenbrock, Julien Y. Dutheil

## Abstract

Meiotic recombination is an important driver of evolution. Variability in the intensity of recombination across chromosomes can affect sequence composition, nucleotide variation and rates of adaptation. In many organisms recombination events are concentrated within short segments termed recombination hotspots. The variation in recombination rate and positions of recombination hotspot can be studied using population genomics data and statistical methods. In this study, we conducted population genomics analyses to address the evolution of recombination in two closely related fungal plant pathogens: the prominent wheat pathogen *Zymoseptoria tritici* and a sister species infecting wild grasses *Zymoseptoria ardabiliae*. We specifically addressed whether recombination landscapes, including hotspot positions, are conserved in the two recently diverged species and if recombination contributes to rapid evolution of pathogenicity traits. We conducted a detailed simulation analysis to assess the performance of methods of recombination rate estimation based on patterns of linkage disequilibrium, in particular in the context of high nucleotide diversity. Our analyses reveal overall high recombination rates, a lack of suppressed recombination in centromeres and significantly lower recombination rates on chromosomes that are known to be accessory. The comparison of the recombination landscapes of the two species reveals a strong correlation of recombination rate at the megabase scale, but little correlation at smaller scales. The recombination landscapes in both pathogen species are dominated by frequent recombination hotspots across the genome including coding regions, suggesting a strong impact of recombination on gene evolution. A significant but small fraction of these hotspots co-localize between the two species, suggesting that hotspots dynamics contribute to the overall pattern of fast evolving recombination in these species.

## Introduction

Meiotic recombination is a fundamental process, which in many eukaryotes shapes genetic variation in populations and drives evolutionary changes. Studies based on experimental and empirical data have demonstrated that recombination in sexual organisms plays a crucial role in defining genome-wide neutral and non-neutral nucleotide variation patterns [1,2], rates of protein evolution [3], transposable elements distribution [4], GC content [5], and codon usage bias [6]. Despite the ubiquitous occurrence of recombination, however, the mechanisms that determine the genome-wide and temporal distribution of crossover events are still poorly understood in most species.

Accurate genome-wide recombination maps are essential for studying the genomics and genetics of recombination. Recombination rates have been recorded in many species by direct observations of meiotic events using genetic crosses or pedigrees (for example [7–9]). Pedigree studies however rely on a large numbers of individuals and produce only low-resolution rate estimates because of the relatively low number of meiotic events that can practically be observed [10]. Furthermore, many microbial eukaryotic species, including important pathogens, are difficult or even impossible to cross under laboratory conditions [11]. While experimental measures of recombination rate can be challenging in many species, advances in statistical analyses provide powerful tools to generate fine-scale recombination maps using population genomic data (e.g., [12–14]). These methods are based on genome-wide patterns of linkage disequilibrium among single nucleotide polymorphisms (SNPs) and have the potential to capture the history of recombination events in a population sample. Thus, recombination studies based on population genomic data have provided detailed insights into the genomics of recombination in a range of species [15–18]. In many organisms, but not all, the majority of recombination events tend to concentrate in short segments termed recombination hotspots [13,19]. In the human genome, more than 25.000 recombination hotspots have been identified, with a number of them showing a more than hundred-fold increase in recombination rates and exhibiting a strong impact on the overall recombination landscape and genome evolution in general [12,15,20].

Comparative analyses of recombination maps between closely related species have shed light on the dynamics of recombination landscapes in different taxa. A comparative analysis of recombination landscapes of chimpanzees and humans found a strong correlation of recombination rates at broad scales (whole chromosome and megabase scale), whereas fine-scale recombination rates were considerably less conserved because of non-overlapping recombination hotspots [21]. The localization of recombination hotspots in primates and mice is in large part determined by PRDM9, a histone methyltransferase with an array of DNA-binding Zn-finger [22]. In some species, including species without PRDM9, such as yeast, plants, birds and some mammals, recombination hotspots associate with particular functional features such as transcription start and stop sites as well as CpG islands [16,18,23–25]. A model developed to explain the association of recombination hotspots and functional elements proposes that a depletion of nucleosome occupancy at these sites increases the accessibility of the recombination machinery [26,27]. Indeed, in the fission yeast *Schizosaccharomyces pombe* and the Brassicaceae plant *Arabidopsis thaliana* meiotic recombination hotspots were shown to co-localize with nucleosome-depleted regions supporting a link between chromatin structure and recombination in these species [27,28].

Although many pathogens and parasites are sexual, the impact of recombination on the evolution of their genomes has been rarely addressed [29]. Genome studies have revealed exceptionally high rates of sequence evolution in some filamentous pathogens including oomycetes and fungi [30,31]. Transposable elements (TEs) play an important role in shaping the architecture and size of these pathogen genomes. TEs have often been found to be enriched in specific genomic compartments such as accessory chromosomes and repeat-rich regions that further encode virulence related genes (reviewed in [31]). Increased mutation rates in TE-rich regions has been shown to contribute to the rapid evolution of new virulence specificities in pathogens [32–34]. While TEs may contribute to the rapid evolution of specific genome compartments, little is known about processes of genome evolution in eukaryotic pathogens. As recombination can be an important driver of overall genome evolution in pathogen species, we here set out to investigate patterns of recombination in plant pathogenic fungi. We focused on the economically important wheat pathogen *Zymoseptoria tritici,* which causes septoria leaf blotch on wheat. *Z. tritici* originated in the Middle East during the Neolithic revolution and has co-evolved and dispersed with its host since early wheat domestication [35]. A close relative of *Z. tritici*, *Zymoseptoria ardabiliae*, has been isolated from wild grass species in the Middle East [36]. The two pathogen species diverged recently but have non-overlapping host ranges and show some differences in morphology and host infection patterns [36,37]. Both species undergo frequent sexual recombination, which result in the formation of ascospores that serve as a mean of long distance wind dispersal and primary infection of new hosts [37]. The co-linear genomes of *Z. tritici* and *Z. ardabiliae* share 90% nucleotide similarity on average, thus providing an excellent resource for comparative analyses of genome evolution [37]. The 40Mb haploid genome of the reference *Z. tritici* isolate comprises 21 chromosomes of which eight are accessory chromosomes [38]. These highly variable chromosomes are characterized by presence/absence variation, structural variation, high repeat content and low gene densities [38,39]. Interestingly, the accessory chromosomes are partly conserved among several species in the genus *Zymoseptoria*, suggesting that these small chromosomes have been maintained over long evolutionary times predating the divergence of species [37].

In a previous study, we applied a whole-genome coalescence approach to generate a map of incomplete lineage sorting of the ancestral species of *Z. tritici* and another closely related species, *Z. pseudotritici* [37]. We found evidence of a high recombination rate in the ancestral species (genome average 46cM/Mb) and showed a significantly higher proportion of sites showing incomplete lineage sorting in regions with high recombination rate. The existence of high recombination rates in the genus *Zymoseptoria* was recently supported by experimental data. Croll and colleagues generated a linkage map of *Z. tritici* from two independent crosses of Swiss field isolates [40]. This map based on actual crossing-over events along the 40Mb genome, confirms the high recombination rates (genome average 66 cM/Mb, measured in windows of 20 kb) in the present-day pathogen species. Interestingly, the study also reported large differences between the two independent crosses of *Z. tritici*, suggesting that the recombination landscape is highly dynamic in this pathogen [40].

In this study we addressed the evolution of recombination rate in fungal pathogens. We applied a population genomics approach to generate a fine-scale recombination map of the two recently diverged species *Z. tritici* and *Z. ardabiliae.* This allowed us to infer and compare fine-scale genome-wide patterns of recombination rates in the two species and investigate the evolution of recombination landscapes. We first of all confirm the exceptionally high recombination rates as also observed in a previous coalescence-based genome analysis and as shown by experimental crosses [37,40]. Furthermore, we identify 2,578 and 862 recombination hotspots in *Z. tritici* and *Z. ardabiliae* respectively. Intriguingly, detailed analyses of the recombination hotspots show not only a comparatively higher hotspot frequency in the wheat pathogen but also the occurrence of stronger hotspots in *Z. tritici*. Our findings confirm that recombination rate landscapes are highly dynamic across time in the two fungal pathogens. Furthermore, the prominence of dynamic recombination hotspots in genes suggests a high impact on gene evolution, a finding that is unprecedented in other species.

## Results and Discussion

### Genome alignments and SNP calling

A total of 30 whole haploid genome sequences was used to infer the recombination landscapes of the two species *Z. tritici* and *Z. ardabiliae*. First, we generated *de novo* genome assemblies of 10 *Z. tritici* and 13 *Z. ardabiliae* isolates previously not studied (Supplemental Table S1). The haploid genomes, including additional three *Z. tritici* and four *Z. ardabiliae* genomes already published [37], were aligned for each species, resulting in multiple genome alignments of 40.8Mb for *Z. tritici* and 32.4Mb for *Z. ardabiliae*.

Recombination analyses rely on single nucleotide polymorphism (SNP) data. However, erroneously called SNPs or alignment errors can greatly bias linkage disequilibrium (LD) inference. To generate high-quality SNP datasets we therefore filtered the genome alignments (see Materials and Methods) to retain only the alignment blocks in which all isolates were represented. This filtering yielded genome alignments of 27.7 and 28.2 Mb for *Z. tritici* and *Z. ardabiliae,* respectively (Table 1). We further filtered the alignments to mask ambiguously aligned positions, leading to a final alignment size of 27.3 Mb for *Z. tritici* and 27.7 Mb for *Z. ardabiliae.* Less than 2% of the final alignment contained repetitive sequences, including transposable elements. In the case of *Z. tritici*, repeat regions have been filtered out during the alignment quality checking, while in the case of *Z. ardabiliae* for which no telomere-to-telomere sequencing is available, most repeats were poorly assembled and therefore virtually absent from the original alignment (Table 1). After filtering, we identified 1.48 million SNPs in *Z. tritici* and 1.07 million SNPs in *Z. ardabiliae*, which correspond to nucleotide diversities measured as Watterson’s θ of 0.0139 in *Z. tritici* and 0.0087 in *Z. ardabiliae* (Table 1). Thus, despite the larger sample size, *Z. ardabiliae* shows a much lower SNP density and sequence diversity than the wheat pathogen *Z. tritici*.

**Table 1:** Summary of genome alignment processing and whole-genome SNP

|  | *Z. tritici* |  | *Z. ardabiliae* |  |
| --- | --- | --- | --- | --- |
| Size of sequenced reference genome | 39,686,251 bp |  | 31,546,591 bp |  |
| Number of exonic sites in reference genome | 17,296,247 bp (43.6%) |  | 15,570,421 bp (49.4%) |  |
| Number of haplotypes | 13 |  | 17 |  |
| Summary genome alignment | Total alignment length | Number of alignment blocks | Total alignment length | Number of alignment blocks |
| MultiZ alignment | 40.8 Mb | 21,500 | 32.4 Mb | 22,296 |
| Splitting in max 10 Kb | 40.8 Mb | 21,904 | 32.4 Mb | 23,001 |
| MAFFT Realignment | 40.5 Mb | 21,904 | 32.2 Mb | 23,001 |
| Keep blocks with all strains | 27.7 Mb | 6,455 | 28.2 Mb | 7,117 |
| Filter 1 | 27.5 Mb | 15,703 | 28.0 Mb | 18,402 |
| Filter 2 | 27.3 Mb | 18,785 | 27.7 Mb | 26,074 |
| Percentage of repeated sequences in initial alignment | 19.74% |  | 3.36% |  |
| Percentage of repeated sequences in final alignment | 0.93% |  | 1.38% |  |
| Total number of SNPs | 1,483,950 |  | 1,069,014 |  |
| Total number of analyzed SNPs (biallelic, no unresolved state) and percent of total SNPs | 1,438,385 (96.9%) |  | 1,035,158 (96.8%) |  |
| Total number of SNPs in exons and percent of total SNPs | 713,733 (48.1%) |  | 403,895 (37.8%) |  |
| Total number of analyzed SNPs in exons (biallelic, no unresolved state), and percent of total analyzed SNPs in exonsTotal number of analyzed SNPs in exons (biallelic, no unresolved state), and percent of total analyzed SNPs in exons | 690,096 (96.7%) |  | 396,247 (98.1%) |  |
| Summary SNP analyses | 1 Mb windows | 100 kb windows | 1 Mb windows | 100 kb windows |
| Min. number of SNPs | 143 | 0 | 0 | 0 |
| Median number of SNPs | 43,680 | 3,556 | 1,598 | 634 |
| Max. number of SNPs | 102,400 | 15,170 | 33,680 | 20,110 |
| Diversity (median of Watterson's theta in windows of 10 kb) | 0.0139 |  | 0.008663 |  |

### Inference of fine-scale recombination maps

We estimated and compared the local recombination rates in *Z. tritici* and *Z. ardabiliae* using two methods implemented in the Ldhat [41] and Ldhelmet [13] packages. Both methods estimate the local population recombination rates based on the LD between SNPs in a given genome dataset using a composite likelihood method. The methods infer the population-scaled recombination rate ρ across the genome, based on an a priori specified population mutation rate θ. The parameter ρ relates to the actual recombination frequency by the equation ρ = 2N_*e*_* *r* for haploid individuals, where N_*e*_ is the effective population size and *r* is the per site rate of recombination across the region. Inferring r from ρ therefore requires knowledge of N_*e*_. Furthermore, in *Zymoseptoria* species, sexual reproduction is not obligatory and may vary from year to year with environmental conditions and the availability of compatible hosts and mating partners, rendering the estimation of r very difficult without any additional knowledge of the amount of clonal reproduction. To avoid the bias of incorrect assumptions we therefore further analysed and compared the recombination maps of *Z. tritici* and *Z. ardabiliae* based on the parameter ρ.

As θ substantially varies along genomes and between species, we generated recombination maps using three scaled effective population size values as inputs (θ = 0.05, 0.005 and 0.0005). For both Ldhat and Ldhelmet, we find that the three different input θ values only have a marginal influence on the recombination rate estimates obtained (Fig. 1A). We therefore proceeded with the recombination map estimated using a θ of 0.005, similar to the median of θ values estimated in 10-kb windows in *Z. tritici* (θ = 0.0139) and in *Z. ardabiliae* (θ = 0.0087) (Table 1).

**Figure 1:**
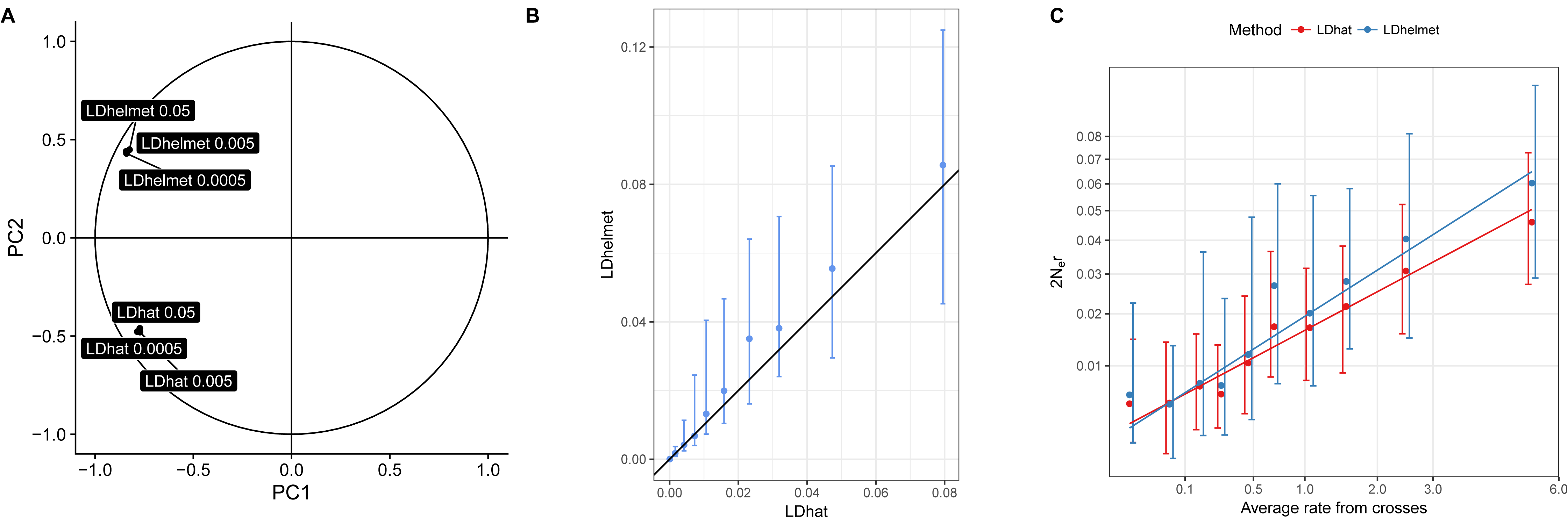
Correlations among recombination maps in *Z. tritici* show highly correlated estimates from two composite likelihood methods. A) Correlation circle of the six population genomic recombination maps based on the two first factors of a principal component analysis. The programs Ldhat interval [41] and Ldhelmet [13] were both used with three distinct input scaled effective population sizes (Θ) of 0.0005, 0.005 and 0.05. B) Correlation of the Ldhat and Ldhelmet maps with Θ = 0.005. The Ldhat map was discretized into 10 categories with equal number of points. The points and error bars represent the median, first and third quartile of the distribution for each category. C) To assess the quality of the inferred recombination maps, genome-wide estimates of recombination were correlated with a genetic map obtained by experimental crossing of *Z. tritici* isolates. Y-axis: Population genomic maps were obtained by Ldhat and Ldhelmet with a scaled population size of 0.005 and x-axis: average recombination map from two independent crosses [40]. Points and error bars represent the median, first and third quartile of the distribution for each category, obtained as in B).

To assess the performance of the two methods and input parameters for the fungal datasets, we first compared the inferred recombination maps of *Z. tritici* with data from previously published genetic maps [40]. We compared both the Ldhat and Ldhelmet recombination maps with the genetic maps created from two sexual crosses of Swiss *Z. tritici* isolates, 3D7x3D1 and SW5xSW39 [40]. The recombination maps estimated by Ldhat and Ldhelmet from SNP data both correlate with the genetic maps confirming that the composite likelihood methods allow us to assess the recombination landscapes in the fungal pathogens (Table 2 and Fig. 1B). We find a significant correlation between the Ldhat map and the two genetic maps (3D7x3D1, Kendall's rank correlation test, τ = 0.27, p-value < 2.2e-16 and SW5xSW39, Kendall's rank correlation test, τ = 0.23, p-value < 2.2.e-16). Using an average recombination rate of the 3D7x3D1 and SW5xSW39 crosses the correlation further increases (Kendall's rank correlation test, τ = 0.29, p-value < 2.2.e-16) (Table 2 and Fig. 1B). While correlated, the new recombination maps of *Z. tritici* encompasses more than 1 million SNPs and thereby provides a considerably finer resolution of the recombination landscape in *Z. tritici* than previously obtained from experimental crosses (based on ca 23,000 SNPs) [40]. The same correlation analyses using the Ldhelmet map show consistent results with slightly lower correlations (Kendall's rank correlation test, τ = 0.24 for the cross 3D7x3D1, and 0.20 for the cross SW5xSW39 and 0.25 using the average of the two crosses; all p-values < 2.2e-16) (Table 2). These correlations, although highly significant, have relatively small size effects. However, it is noteworthy that also the correlation between the two Swiss crosses 3D7x3D1 and SW5xSW39 only is 0.43 (Kendall’s rank correlation test, p-value < 2.2e-16) supporting a high variability in recombination even between individual crosses of *Z. tritici* (Table 2). Based on the comparison of the outputs of Ldhat and Ldhelmet, we decided to use the Ldhat map as our reference population map for the remaining of this study. We next investigated the impact of possible confounding factors on the recombination rate estimates including SNP densities, possible sequencing errors, population structure and natural selection.

**Table 2:** Robustness of the population recombination map and correlation with cross-over maps

| Data | Map | 3D7x3D1 |  |  | SW5xSW39 |  |  | Average |  |  |
| --- | --- | --- | --- | --- | --- | --- | --- | --- | --- | --- |
|  |  | Correlation | P-value | CI | Correlation | P-value | CI | Correlation | P-value | CI |
| Unfiltered | LDhat | 0.27 | 2.22E-60 | [0.24,0.30] | 0.23 | 5.11E-44 | [0.20,0.26] | 0.29 | 4.61E-69 | [0.26,0.32] |
|  | LDhelmet | 0.24 | 1.68E-47 | [0.21,0.27] | 0.20 | 8.18E-33 | [0.17,0.23] | 0.25 | 1.11E-52 | [0.22,0.28] |
|  | Average | 0.26 | 2.59E-56 | [0.23,0.29] | 0.22 | 9.81E-40 | [0.19,0.25] | 0.27 | 4.71E-63 | [0.24,0.30] |
|  | LDhat intergenic | 0.20 | 9.54E-32 | [0.17,0.24] | 0.18 | 9.54E-25 | [0.15,0.21] | 0.22 | 2.65E-37 | [0.19,0.25] |
|  | LDhelmet intergenic | 0.22 | 9.73E-37 | [0.19,0.25] | 0.19 | 1.89E-28 | [0.16,0.23] | 0.23 | 1.79E-42 | [0.20,0.27] |
|  | LDhat no singleton | 0.21 | 1.18E-36 | [0.18,0.24] | 0.17 | 1.81E-24 | [0.14,0.20] | 0.21 | 6.88E-39 | [0.18,0.24] |
|  | LDhat no structure | 0.25 | 7.77E-54 | [0.22,0.29] | 0.23 | 7.90E-43 | [0.19,0.26] | 0.26 | 1.48E-59 | [0.23,0.29] |
| Filtered | LDhat | 0.31 | 3.36E-76 | [0.27,0.33] | 0.28 | 1.94E-64 | [0.25,0.31] | 0.34 | 2.10E-96 | [0.31,0.37] |
|  | LDhelmet | 0.26 | 4.45E-57 | [0.23,0.29] | 0.22 | 3.35E-40 | [0.19,0.25] | 0.28 | 1.34E-64 | [0.25,0.31] |
|  | Average | 0.29 | 9.54E-69 | [0.26,0.32] | 0.25 | 2.62E-51 | [0.22,0.28] | 0.31 | 5.65E-80 | [0.28,0.34] |
|  | LDhat intergenic | 0.20 | 5.39E-30 | [0.17,0.23] | 0.18 | 3.70E-24 | [0.15,0.21] | 0.22 | 1.90E-36 | [0.19,0.25] |
|  | LDhelmet intergenic | 0.23 | 1.81E-37 | [0.19,0.26] | 0.19 | 5.06E-26 | [0.15,0.22] | 0.24 | 6.40E-42 | [0.20,0.27] |
|  | LDhat no singleton | 0.29 | 6.92E-66 | [0.25,0.32] | 0.25 | 6.22E-52 | [0.22,0.28] | 0.31 | 8.55E-79 | [0.28,0.34] |
|  | LDhat no structure | 0.29 | 1.24E-71 | [0.26,0.32] | 0.29 | 2.58E-67 | [0.26,0.32] | 0.32 | 3.66E-88 | [0.29,0.35] |

Note: Correlation values are Kendall's tau. CI: 95% confidence interval obtained by 10,000
bootstraps. SW5xSW39 and 3D7x3D1 correspond to crosses in [40]..

### SNP density and filtering based on confidence intervals

Ldhat and Ldhelmet have been developed for recombination analyses in animals [13,21,41] and their performance on data from haploid eukaryotes with high recombination rates have not been tested. Therefore, we next assessed the robustness of the composite likelihood approach using simulations with distinct sample sizes and SNP densities. We report that the *interval* program infers recombination rate with the highest reliability for intermediate diversity levels (θ = 0.0005 or 0.005). Furthermore, while larger sample size decrease the variance in estimate, we show that Ldhat reliably infers recombination when as few as 10 haploid genomes are used (Fig. 2). We observe that ρ generally tends to be underestimated and its estimation variance is larger for small sample sizes. Yet better estimates can be obtained by discarding all estimates with a 95% confidence interval at least equal to two times the mean. Interestingly, this filtering has the strongest effect for highly diverse regions (θ = 0.05), where the raw estimates of Ldhat appear to be highly underestimated even for large sample sizes (n = 100). Discarding estimates with large confidence intervals efficiently suppress this bias (Fig. 2). We also note that the inference bias is stronger for low recombination rates, and that this effect is independent of the sample size (Fig. 2). Based on these simulation results, we similarly filtered our recombination estimates based on the 95% confidence interval reported by Ldhat. This filtering discards 49% and 20% of all SNP pairs for *Z. tritici* and *Z. ardabiliae*, respectively. The large difference between the two data sets is imputable to the much higher nucleotide diversity of *Z. tritici*. When compared with the genetic map [40], the filtered map of *Z. tritici* shows a correlation of 0.34 (Kendall’s rank correlation test, p-value < 2.2e-16, Table 2). Interestingly, correlations between the genetic map and the linkage disequilibrium (LD) map inferred with Ldhat increases with increased window size: using 500 kb windows, the correlation becomes 0.43 (Kendall’s tau, p-value = 0.000206).

**Figure 2:**
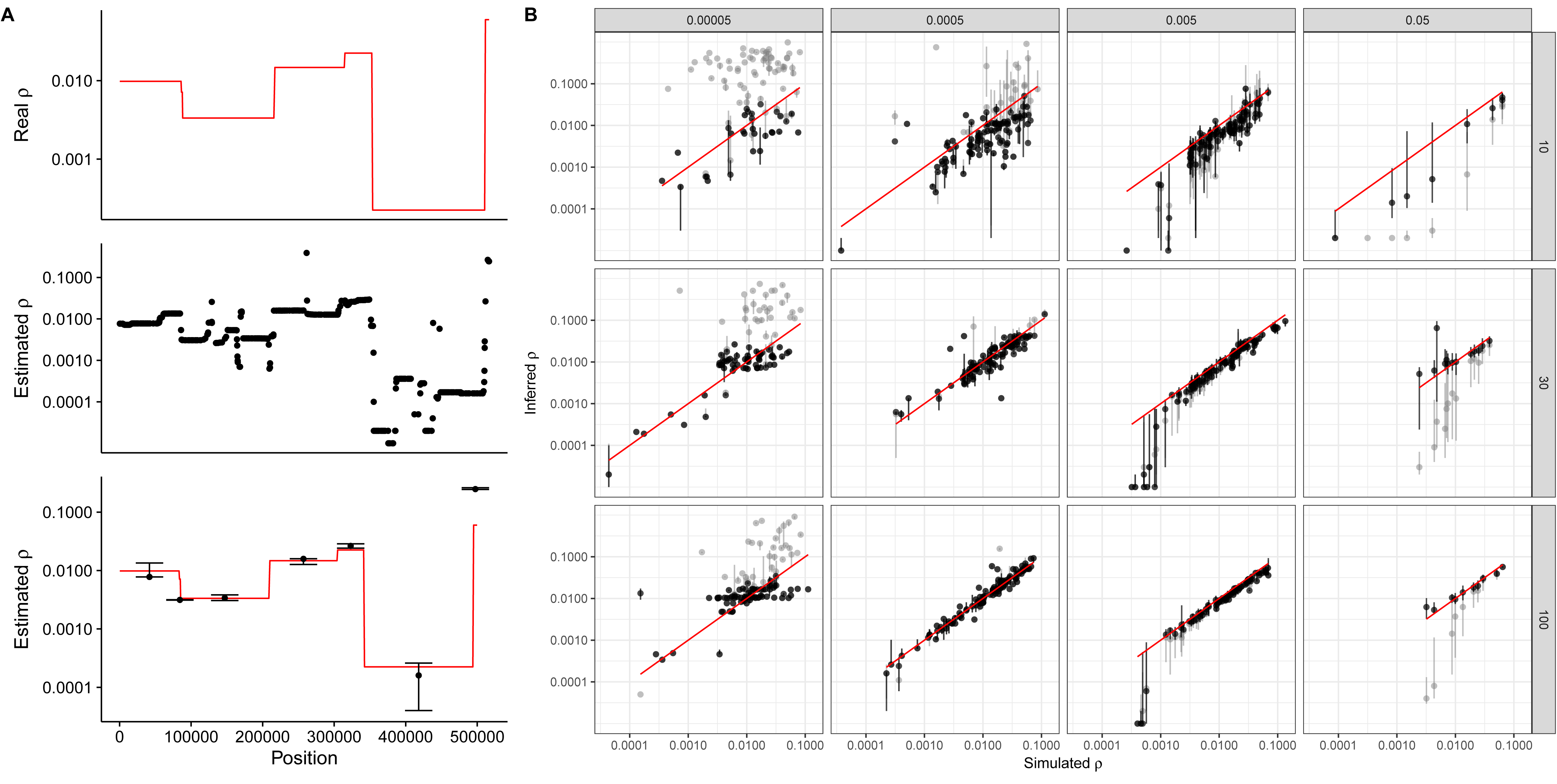
effect of sample size and diversity on the estimation of recombination rate by Ldhat. 10 Mb regions (1Mb for regions with θ = 0.05) were simulated using a coalescent model with variable recombination rate: random segments were generated by sampling lengths from an exponential distribution, and rates from the observed distribution of recombination rates. A) Example of a 500 kb region, with variable recombination rate (red line), LDhat estimates between pairs of SNPs (middle panel) and median (with first and third quartiles as error bars) for each segment of uniform recombination (bottom panel). B) Inferred vs. true recombination rate for different nucleotide diversity values (θ = 2 Ne u) and sample sizes. Each dot corresponds to a region with constant recombination rate in the simulated alignment, as shown in A). Bars indicate the 1^st^ and 3^rd^ quartiles of Ldhat estimates for the region. Grey points are raw estimates; black points are computed from filtered estimates (see Methods). The red diagonal line shows the 1:1 ratio. Columns indicate distinct population mutation rates and rows distinct sample sizes (number of haploid genomes).

### Putative sequencing errors

Sequencing errors can impact linkage disequilibrium estimates as they appear as unlinked singletons in population genomic datasets. Such potential effects are partially accounted for at two levels in our analyses. First, our data sets are based on *de novo* assembly of each individual genomes, which already corrects for putative sequencing errors in the sequencing read outs. Second, many of the SNPs discarded for having a large confidence interval in the estimation of recombination rates by LDhat are singletons. In order to further assess the potential impact of putative sequencing errors, we ran LDhat on the *Z. tritici* data set after discarding all singletons and filtering for confidence interval as described above. The resulting recombination map appeared to be highly correlated to the map including all singletons (Table 2 and Figure S1), and the correlation of the filtered map with the crossing-over map was significant, but less than when including them (Kendall’s tau = 0.3083, p-value < 2.2E-16) (Table 2). We therefore conclude that sequencing errors have no significant impact on our inferred recombination maps.

### Effect of population structure

Previous studies reported that *Z. tritici* strains are sampled from a globally panmictic population [42,43]. However in a recent study based on whole genome data, we report evidence for slight population structure, notably between Iranian isolates vs. European isolates [44]. In order to assess whether this structure could bias our recombination estimates, we generated a new recombination map using LDhat on a reduced sample of eight strains of *Z. tritici*. We excluded the two Iranian isolates in our dataset as well as three German isolates forming a separate cluster, yet non-significant. We report that the resulting map is highly correlated to our recombination map (Figure S1) as well as to the crossing-over map (Kendall’s tau = 0.3237, p-value < 2.2E-16) (Table 2) suggesting that population structure has little impact on our inference of recombination rate. Because the resulting correlation with the crossing-over map was slightly lower than when using the complete data set, we use the complete dataset for the further analyses.

As little is known about the population structure of *Z. ardabiliae*, we conducted additional simulations in order to assess the putative impact of structure on the inference of recombination rate. We used a five-islands structure model, with sample sizes equal to 2, 3, 4, 5 and 6 in each deme, respectively, with a total sample size of 20, comparable to the 17 genomes of the *Z. ardabiliae* data set analyzed here. Migration rates between demes were symmetrical and all equal, and we tested several rates. We find that while ρ is systematically overestimated in the presence of population structure, it is remarkably proportional to the true value, in particular after filtering on the confidence intervals (Figure S2). Population structure, if any, is therefore not expected to bias our comparison of recombination rates along the genome. In addition, these results suggest that the true recombination rate in *Z. ardabiliae* is potentially even lower than the value reported here.

## Coding sequences

Recombination inference based on patterns of linkage disequilibrium is affected by various patterns of selection. The genomes of *Z. tritici* and *Z. ardabiliae* are gene dense and protein-coding genes occupy nearly 50% of the sequences. We therefore considered the impact of selection on our recombination inference in the two species assuming lower selection in non-coding regions. To this end, we compared the previously published genetic map with estimates of ρ exclusively in the intergenic regions, excluding coding sequences and 500-bp up and downstream of the annotated genes (Table 2 and Figure S1). These analyses based on non-coding sequences and filtering of SNPs based on the confidence interval of recombination rate estimates resulted in correlations of 0.22 for the Ldhat map and the average of the two genetic crosses (Kendall’s rank correlation test, p-value < 2.2e-16) and for the Ldhelmet map (Kendall’s rank correlation test, p-value < 2.2e-16). Thus, the best correlations of LD based on the recombination maps and genetic crosses are obtained when coding regions are included (Table 2). The finding suggests that the composite likelihood method provides robust estimates of recombination, even in regions likely to deviate from purely neutral evolution. Based on these simulation results, we chose to use the Ldhat-inferred recombination rates on the full genome, with an input θ = 0.005 and filtered according to confidence intervals, for both *Z. tritici* and *Z. ardabiliae*.

### A five fold higher population scaled recombination rate in *Z. tritici*

The inference of ρ across the genomes of *Z. tritici* and *Z. ardabiliae* reveals highly heterogeneous recombination landscapes in both species (Fig. 3 and Supplementary Data 1). We find a five-fold higher recombination rate in *Z. tritici* than in *Z. ardabiliae*: the mean values of ρ are 0.0217 and 0.0045 for *Z. tritici* and *Z. ardabiliae*, respectively. As ρ = 2N_e_\**r*, where *r* is the actual recombination rate per generation per nucleotide and N_e_ is the effective population size, this five-fold difference might reflect differences in *r*, or global differences in N_e_. Furthermore, the inferred parameter ρ reflects the historical rates of recombination in the two species, which may have varied according to different demographic events since their divergence. Nonetheless, the nucleotide diversity estimated by Watterson’s θ, is 1.6 times higher in *Z. tritici* than in *Z. ardabiliae*, indicating that different population sizes alone cannot explain the observed difference in recombination rates assuming that the two species have comparable mutation rates. The higher value of ρ estimated in *Z. tritici* thus likely reflects a higher actual recombination rate, in the past or presently, in the wheat pathogen compared to *Z. ardabiliae*.

**Figure 3:**
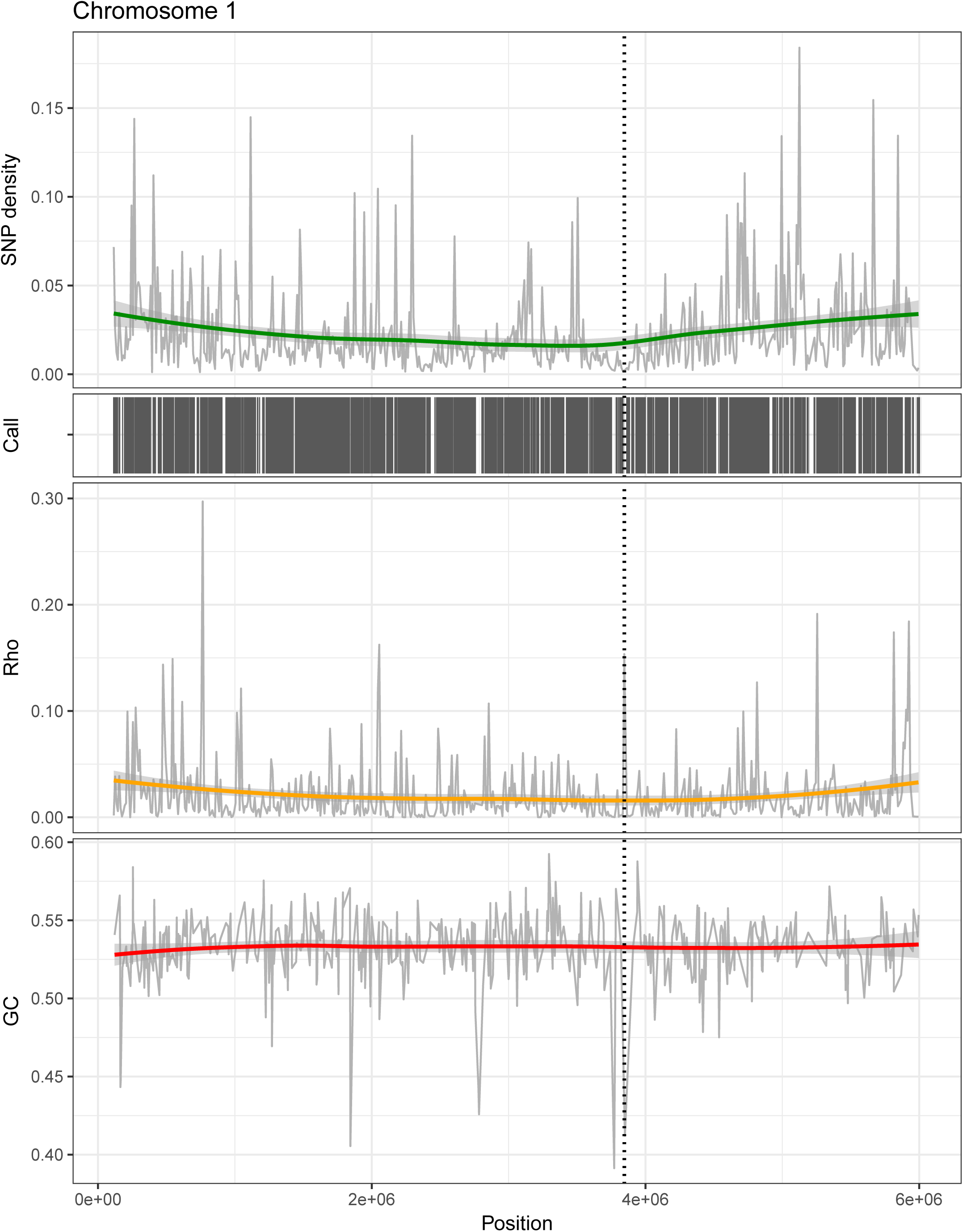
Variation in recombination rate across chromosomes. Based on the population genomics data of *Z. tritici* and *Z. ardabiliae*, genome-wide patterns of recombination are estimated. Patterns of variation across chromosome 1 of *Z. tritici* is shown as example. A) SNP density in 10 kb windows with corresponding smoothing curve. B) Distribution of called sites along the chromosome in black, corresponding to the regions that were included in the analyses. C) Estimates of the population recombination rate ρ show a highly heterogeneous small-scale recombination landscape across the chromosomes. D) Observed GC content. The position of the centromere is marked over the chromosome plots as a vertical stippled line.

### Recombination on small arms of acro-centric chromosomes

Physical factors, such as chromosome length, centromere position or distance to the centromere have been reported to impact broad-scale recombination patterns in eukaryotes [45]. To investigate the rate and distribution of crossover events along the genomes of the two *Zymoseptoria* species, we correlated the inferred recombination maps with features of the well-characterized karyotype of *Z. tritici*. The reference genome sequence of *Z. tritici* consists of 21 fully sequenced chromosomes, including eight so-called accessory chromosomes showing presence/absence polymorphisms between individuals [38]. Furthermore, the exact positions of the centromeres for all chromosomes have been characterized experimentally using a chromatin immunoprecipitation assay targeting the centromere specific protein CenH3 [46]. An interesting finding is that the chromosomes in *Z. tritici* are either acro-centric or near-acrocentric, and every chromosome consequently consists of one long and one short chromosome arm [46]. Because a complete chromosome assembly is not available for *Z. ardabiliae*, we mapped the recombination estimates of *Z. ardabiliae* on the genome of *Z. tritici* to assess the impact of the karyotype structure on recombination rate variation. Similar to findings from other species [45,47], we observe a negative correlation between recombination rate and the size of the thirteen core chromosomes (Kendall’s τ = −0.59 with p-value = 4.29e-3 for *Z. tritici* and τ = −0.72 with p-value = 2.84e-4 for *Z. ardabiliae*; Fig. 4A). This pattern is generally explained by the necessity of one crossing over to occur per chromosome or chromosome arm per generation, resulting in a higher recombination rate on smaller chromosomes (e.g., [24,48]). The significant correlation of the recombination map of *Z. ardabiliae* with the genome structure of *Z. tritici* is an indication of a conserved karyotype of the ancestral species of *Z. tritici* and *Z. ardabiliae*.

**Figure 4:**
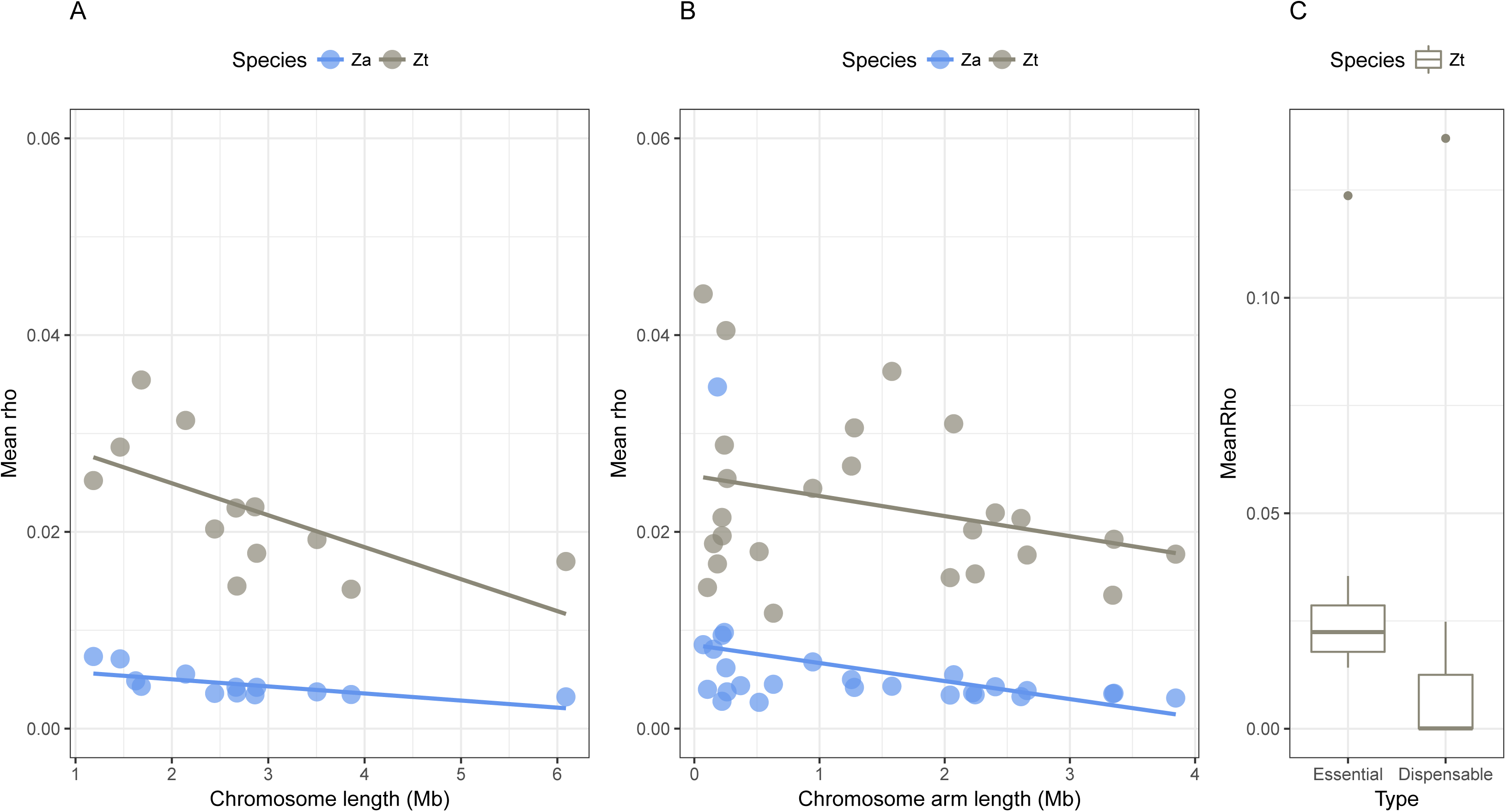
Broad-scale recombination rates in *Z. tritici* and *Z. ardabiliae.* Broad-scaled patterns of recombination rate in *Z. tritici* and *Z. ardabiliae* demonstrate a strong effect of chromosome length and type. A) Mean recombination rate in *Z. tritici* and *Z. ardabiliae* per essential chromosome as a function of the chromosome length. B) Mean recombination rate per essential chromosome arm as a function of the arm length. C) Distribution of mean recombination rate per chromosome in *Z. tritici* as a function of type (essential or accessory).

Given the acro-centric nature of the *Z. tritici* chromosomes we considered to which extent recombination also occurs on the short chromosome arms. If meiosis involves one crossover event per chromosome, then the recombination rate should be correlated with the chromosome size and not the chromosome arm length. However, if meiosis involves one crossover event per chromosome arm, then a higher frequency of recombination should occur on shorter chromosome arms. Correlations between recombination rates and chromosome arm lengths also show negative values, yet only significant in *Z. ardabiliae* (Kendall’s τ = −0.14 with p-value = 0.3356 for *Z. tritici* and τ = −0.42 with p-value = 2.16e-3 for *Z. ardabiliae,* Fig. 4B). The negative correlation observed at the chromosome arm level suggests that meiosis in the *Zymoseptoria* pathogens requires at least one crossing over per chromosome arm and that the small chromosome arms consequently also recombine. The weaker correlations and lack of significance in *Z. tritici* could be due to a fast evolution of centromere positions, erasing the signal of arm-specific recombination rates.

### Extremely weak or absent GC-biased gene conversion in *Z. tritici* and *Z. ardabiliae*

In many species recombination strongly impacts evolution of GC content by a mechanism called GC biased gene conversion (gBGC) [49,50]. The effect of gBGC has been demonstrated in mammals [49,51], birds [52], plants [53] and even bacteria [54]. However, gBGC has never been assessed in fungal species beyond the yeast model, which represents one of the rare organisms for which gBGC was experimentally demonstrated [55]. To study the possible occurrence and impact of gBGC in the *Z. tritici* and *Z. ardabiliae* genomes, we studied the patterns of GC content along the genomes of the two species. We fitted a non-homogeneous, non-stationary model of substitution in 10 kb windows in intergenic regions allowing us to estimate the equilibrium GC content (frequency of GC towards which the sequences evolve) in the extant species. We inferred the dynamics of GC content by comparing the actual GC content of the sequence (observed GC content) with the equilibrium GC content [56]. We find that both the observed and equilibrium GC are highly correlated between *Z. tritici* and *Z. ardabiliae* (Supplemental Fig. 3, Kendall’s rank correlation test, τ = 0.69 and 0.45, p-values < 2.2E-16 for the observed and equilibrium GC content, respectively, essential chromosomes only). However, although both species show similar observed GC content (median of 53.3% for *Z. tritici* and 53.6% for *Z. ardabiliae*) they also show contrasting patterns, with the GC content found to be slightly increasing in *Z. ardabiliae* (median equilibrium GC content on autosomes of 53.8, significantly higher that the observed GC content, Wilcoxon paired rank test, p-value = 0.04712) while decreasing in *Z. tritici* (median equilibrium GC content of 51.6%, which is significantly lower than the observed GC content, Wilcoxon paired rank test, p-value = 2.728e-15).

To assess the impact of recombination on GC evolution we correlated the equilibrium GC content in *Z. tritici* and *Z. ardabiliae* to the recombination maps in the two species. We find overall negative yet weakly or non-significant correlations between GC content and recombination rate (Supplemental Fig. S3), both for observed (Kendal’s tau = −0.047, p-value

= 0.04304 for *Z. tritici* and tau = −0.054, p-value = 0.02253 for *Z. ardabiliae*) and equilibrium GC content (Kendal’s tau = −0.02, p-value = 0.5082 for *Z. tritici* and tau = 0.01, p-value = 0.7128 for *Z. ardabiliae*). These results do not support GC-biased gene conversion as a major mechanism shaping GC content in the two fungal pathogen genomes. To test whether this conclusion could be an artifact of recombination rates estimated from population data, we also correlated the equilibrium GC content with the two previously published genetic maps [40]. Consistent with our finding from the Ldhat-based recombination map, we confirm an absence of correlation between the equilibrium GC content and the crossing-over rate and GC content in *Z. tritici*, (Kendall’s rank test, τ = 0.006 and p-value = 0.7035 for observed GC and τ = −0.024, p-value = 0.1149 for equilibrium GC content).

The absence of correlation between GC content and recombination could also be due to a lack of statistical power due to the overall very homogeneous GC content and recombination large-scale landscapes (recall Figure 3), and the notable absence of isochores that characterize genome composition in other organisms, e.g. in Mammals [57]. As a complementary line of evidence, we investigated the segregation patterns of AT and GC alleles at AT/GC biallelic sites in intergenic regions of both *Z. tritici* and *Z. ardabiliae*, as gBGC is expected to increase the frequency of GC-alleles [58]. We find that the frequency of GC alleles is virtually identical in *Z. tritici* and only slightly higher in *Z. ardabiliae* to the frequency of AT alleles (Table 3), supporting an absence or only weak effect of GC-biased gene conversion in *Z. tritici* and *Z. ardabiliae*, respectively.

**Table 3:** Segregation patterns at AT/GC biallelic sites.

| Species | Frequency of GC alleles | Number of alleles with GC > 50% | Number of alleles with GC < 50% | Ratio GC / (AT + GC) |
| --- | --- | --- | --- | --- |
| *Z. tritici* | 50.41% | 274,517 | 268,589 | 50.55% |
| *Z. ardabiliae* | 51.86% | 261,777 | 236,232 | 52.56% |

### No suppression of recombination in centromeres

Recombination is normally found to be absent in centromeric regions where spindles attach during chromosome segregation (see review by [19]). A known exception is *Drosophila mauritiana*, which, in contrast to *Drosophila melanogaster* and *Drosophila simulans*, shows no suppression of recombination in centromeres [59]. The centromeres of core and accessory chromosomes in *Z. tritici* range from 5.5 kb to 14 kb in size and do not locate in AT rich regions [46] as is otherwise observed for centromeres of other species such as *Neurospora crassa* [60]. Correlating the recombination map of *Z. tritici* with centromere positions, we observe, as in *D. mauritiana*, no significant suppression in recombination rate across the centromeric chromosome regions (Wilcoxon signed rank test on 11 chromosomes for which recombination rate in the centromeric region could be inferred, p-value = 0.5771) (Table 4, Fig. 3). The centromeres of *Z. tritici* exhibit several features common to neocentromeres such as a short length (approx. 10,000 bp in length), lack of enriched repetitive DNA and weakly transcribed genes [46]. We hypothesize that recombination in centromeric sequences has additional implications for evolution of the centromeres in these fungi. A more detailed characterization of chromosome structures and centromere locations in *Z. ardabiliae* is necessary to better understand karyotype evolution in these grass pathogens.

**Table 4:** Recombination and repeat content in centromeres of *Z. tritici*.

|  | Chromosome | Start | Stop | Length | Mean rho | Nb. of SNPs | Mean rho for Full chromosome | Repeat density | TE density |
| --- | --- | --- | --- | --- | --- | --- | --- | --- | --- |
| Essential | 1 | 3839299 | 3851749 | 12450 | 0.229 | 20 | 0.021 | 0.94% | 31.33% |
|  | 2 | 512901 | 521916 | 9015 | 0.053 | 77 | 0.024 | 0.00% | 32.39% |
|  | 3 | 3348307 | 3356535 | 8228 | 0.097 | 269 | 0.025 | 0.00% | 0.00% |
|  | 4 | 217113 | 226545 | 9432 | 0.033 | 421 | 0.028 | 0.00% | 9.88% |
|  | 5 | 2604117 | 2614736 | 10619 | 0.104 | 47 | 0.027 | 0.94% | 28.19% |
|  | 6 | 625186 | 637601 | 12415 | NA | 0 | 0.026 | 3.10% | 37.46% |
|  | 7 | 255824 | 266207 | 10383 | 0.006 | 79 | 0.044 | 0.32% | 0.00% |
|  | 8 | 213892 | 227444 | 13552 | 0.059 | 62 | 0.029 | 0.45% | 39.99% |
|  | 9 | 2067589 | 2076063 | 8474 | 0.015 | 106 | 0.040 | 0.50% | 0.00% |
|  | 10 | 99716 | 109365 | 9649 | 0.016 | 77 | 0.049 | 0.00% | 15.32% |
|  | 11 | 365130 | 373557 | 8427 | NA | 0 | 0.049 | 0.00% | 46.30% |
|  | 12 | 180233 | 188209 | 7976 | 0.001 | 150 | 0.052 | 2.48% | 7.10% |
|  | 13 | 236993 | 242558 | 5565 | 0.015 | 156 | 0.037 | 0.50% | 0.00% |
| Dispensable | 14 | 59960 | 70870 | 10910 | 0.000 | 785 | 0.000 | 0.00% | 35.86% |
|  | 15 | 382500 | 394754 | 12254 | 0.001 | 1098 | 0.001 | 0.86% | 20.04% |
|  | 16 | 332004 | 342592 | 10588 | 0.099 | 83 | 0.023 | 0.00% | 35.97% |
|  | 17 | 406958 | 418893 | 11935 | NA | 0 | 0.000 | 0.24% | 46.85% |
|  | 18 | 159000 | 171999 | 12999 | NA | 0 | 0.159 | 0.00% | 46.62% |
|  | 19 | 148227 | 159387 | 11160 | 0.001 | 4 | 0.000 | 0.76% | 1.38% |
|  | 20 | 94677 | 105169 | 10492 | NA | 0 | 0.008 | 0.30% | 11.86% |
|  | 21 | 340264 | 346657 | 6393 | NA | 0 | NA | 0.31% | 2.33% |

### Absence of recombination on accessory chromosomes

The small accessory chromosomes have previously been well characterized in *Z. tritici* [38]. They differ considerably from the core chromosomes as they display a higher repeat content, lower gene density, overall lower transcription rate and are enriched with different chromatin modifications [39,46,61,62]. Electrophoretic separation of accessory chromosomes from several isolates of *Z. ardabiliae* has shown that this species also comprises accessory chromosomes [37]. In this study we used sequence homology to define the accessory components of the *Z. ardabiliae* genome. We find that the aligned fragments of the accessory chromosomes show very low recombination rates in both species (median ρ = 0.0059 in *Z. tritici* and median ρ = 0.0001 in *Z. ardabiliae* over 13 10-kb windows where both genomes could be aligned, which is 25% and 2% of the autosomal rates, respectively) (Fig. 4C). The lower recombination rates reflect the lower effective population size of accessory chromosomes that are present at lower frequencies in populations of *Z. tritici* and *Z. ardabiliae* compared to the core chromosomes. Furthermore we speculate that frequent structural rearrangements on accessory chromosomes can prevent homologous chromosomes pairings and also contribute to the low recombination rates. Our findings add further evidence to support different evolutionary modes of the two sets of chromosomes (core and accessory chromosomes) contained in the same genome. Suppression of recombination is also found on mating-type chromosomes in other fungi including species of *Neurospora* and *Microbotryum [63–65]*. These regions are characterized by an increased accumulation of transposable elements and structural variants as well as non-adaptive mutations in coding sequences as a consequence of suppressed recombination [65–67].

We also observe a remarkable drop in the recombination rate on the right arm of chromosome 7 (Supplemental Data 1). The right arm of chromosome 7 displays several similarities to the DNA of the accessory chromosomes, including a lower gene density, higher repeat content and less gene transcription [39]. Furthermore, the entire chromosome arm is enriched with the heterochromatic mark H3K27me3, which is similarly enriched on the accessory chromosomes [46]. We previously proposed that this particular chromosome region represents a recent translocation of an accessory chromosome to a core chromosome [46]. This hypothesis is consistent with the observation that the recombination rate of the chromosome arm resembles the overall reduced recombination rate of the accessory chromosomes (Supplemental Data 1).

### High recombination rates in coding sequences of *Z. tritici*

In primates and birds, recombination increases at CpG islands and around transcription start and end sites [16,21,24]. In honeybee recombination rates in introns and intergenic regions are significantly higher than recombination rates in 3’ and 5’ UTRs and coding sequences [68]. It has been proposed that altered chromatin structures such as destabilized nucleosome occupancy at CpG islands and promoters contribute to this fine-scale variation in recombination rate [69]. To determine whether specific sequence features in the fungal pathogen genomes similarly affect the overall recombination landscape, we inferred and compared the mean recombination rates in exons, introns, intergenic regions, and 5’ and 3’ flanking regions (500-bp upstream and downstream CDS regions, respectively) with a minimum of 3 filtered SNPs (Fig. 5A). Overall, we observe significant differences but with small size effects in fine-scale rates of recombination across different genome regions (Kruskal-Wallis test with post-hoc comparisons, FDR set to 1%). In both *Z. tritici* and *Z. ardabiliae* we find the lowest recombination rates in introns and the highest rates in intergenic sequences (Fig. 5A). A lower value of ρ = 2N_e_*r* can result from a reduced N_e_, a reduced *r* or both. N_e_ in the proximity of genes is expected to be lower due to the presence of background selection [70–72]. The highly similar observed recombination rates in coding and non-coding sequences in *Z. tritici* and *Z. ardabiliae* suggests that *r* is not suppressed in these regions in the same way as observed in other organisms. The pattern indicates that other mechanisms define fine-scale recombination rates in these fungi leading to high recombination frequencies in protein-coding sequences.

**Figure 5:**
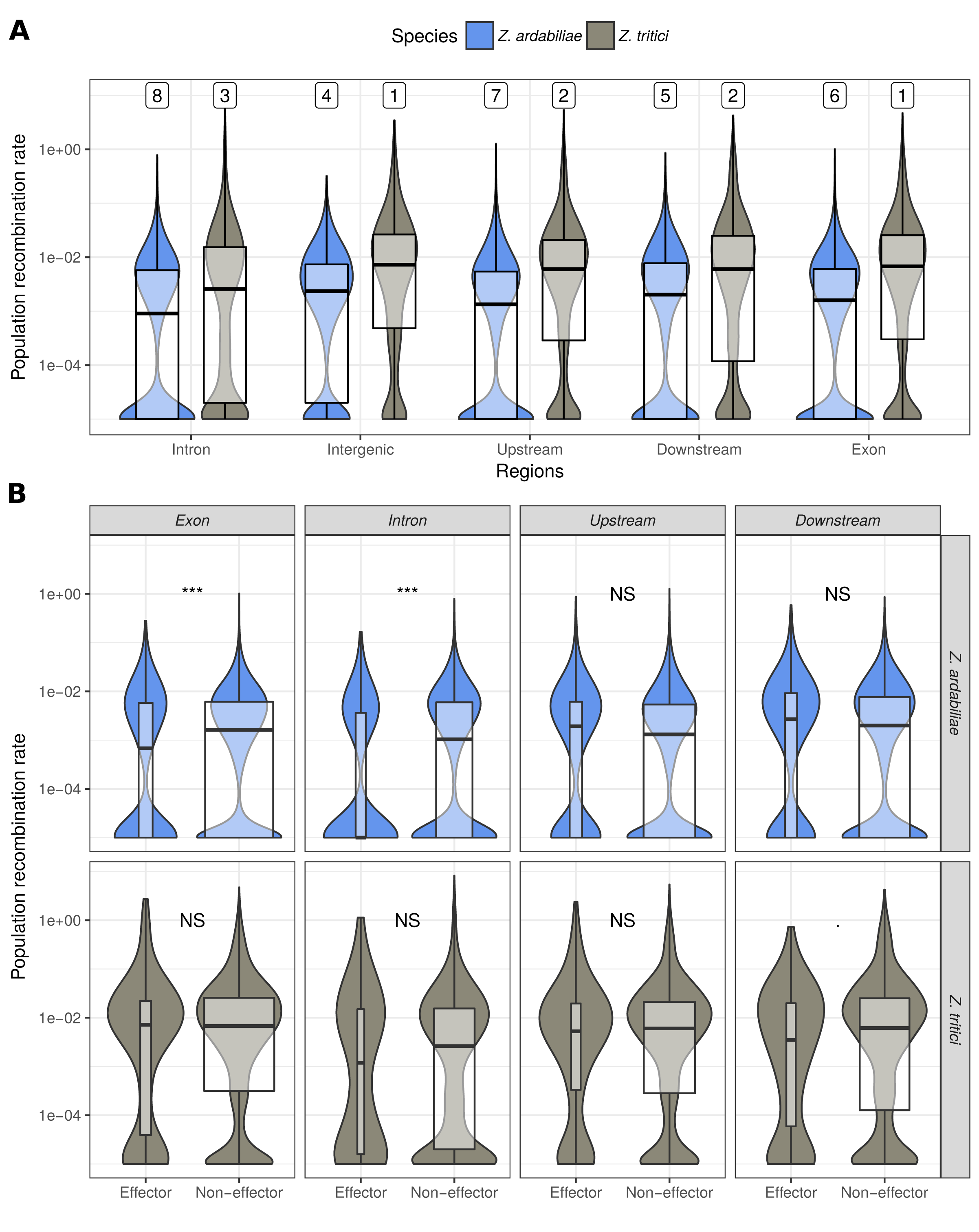
Fine-scale recombination patterns within chromosomes. A) The distribution of recombination rate estimates in different sequence features in *Z. tritici* and *Z. ardabiliae* reveals small, but significant differences among the non-coding, coding and UTR sequences in both species. Top line numbers indicate significance groups by decreasing value of recombination rate. Categories with identical numbers are not significantly different at the 1% level. B) Distribution of recombination rate estimates in exons, introns and UTRs of effector and non-effector genes is shown. Bow widths are proportional to the sample sizes. For *Z. ardabiliae*, the recombination rate in exons and introns is significantly lower in effector genes compared to non-effector genes (Wilcoxon rank test corrected for multiple testing, NS: non significant, *: 5% level, ***: below 0.1% level).

Because of the relatively high rates of recombination in exons of *Z. tritici* and *Z. ardabiliae*, we sought to determine whether recombination could play a particular role in plant-pathogen co-evolution. Plant pathogens interfere with host defenses and manipulate the host metabolism by the secretion of so-called effector proteins produced to target molecules from the host [73]. Antagonistic co-evolution of these interacting proteins is often reflected in accelerated evolution and signatures of positives selection [74]. To assess the role of recombination on effector evolution, we first predicted effector proteins computationally in the secretomes of both species using the EffectorP software [75]. This approach identified 868 putative effector proteins in *Z. tritici* and 1,122 and *Z. ardabiliae*.

By comparing the recombination rates in different genomic regions encoding effector and non-effector genes, we show a significantly lower recombination rate in exons and introns of effector proteins in *Z. ardabiliae* (Wilcoxon rank test, p-value = 1.305e-4 (exons) and 2.534e-5 (introns), p-values corrected for multiple testing) (Fig. 5B). The differences are mostly driven by an excess of zero estimates in effector-encoding regions in *Z. ardabiliae*, as visible on the distribution of measures (Fig. 5B). Discarding these regions with a mean recombination of zero leads to non-significant differences between effector and non-effector genes. A recombination rate estimated to zero can either be due to suppression of recombination in the region or to an estimation error. Intron and exons with a recombination estimate of zero in *Z. ardabiliae* are found to be shorter and to have a higher SNP density (Supplementary Data 3). While these differences are significant, they are of a small size and are unlikely to be a cause of estimation error, and the suppression of recombination in some effector genes of *Z. tritici* therefore appears as a biological signal which origin remains to be elucidated by detailed analysis of these regions.

### Large scale but not fine scale correlation of recombination landscapes in *Z. tritici* and *Z. ardabiliae*

Recombination landscapes have been compared in different model species to assess the extent of conservation of recombination rate variation. Broad-scale recombination rates in zebra finches and long-tailed finches have similar levels and present correlation factors as high as 0.82 and 0.86 at the 10-kb and 1-Mb scales, respectively [16]. Similarly, broad-scale recombination rates in human and chimpanzee tend to be conserved with few exceptions such as the human chromosome 2, which originates from a chromosome fusion in the human lineage [21]. However, when comparing the recombination rates of more distantly related mammal species, the correlation of recombination rates decreases even when comparing homologous syntenic blocks [45]. In studies of mammals and fruit flies, it is considered that the recombination landscape evolves as a result of evolution of other sequence variables [45], and the dynamics of fine-scale recombination rates including the positions of hotspots [13,15].

To address the evolution of recombination landscapes in *Z. tritici* and *Z. ardabiliae* we compared the genome-wide recombination maps of the two species. We previously reported that the genomes of the two species show a high extent of co-linearity and we found a mean sequence divergence of d_*xy*_= 0.13 substitutions per site [37]. Here, we first aligned the two reference genomes of *Z. tritici* and *Z. ardabiliae* to compare recombination rates in homologous genome regions (Fig. 6, see Materials and Methods). Next, we calculated the average recombination rate in non-overlapping windows with at least 100 SNPs in each species, which resulted in 3,851 windows for which recombination in both species could be averaged. The two maps show a moderate yet highly significant correlation (Kendall's rank correlation test, τ = 0.2327, p-value < 2.2e-16, Fig. 7A), which suggests certain similarities in the recombination landscape of the two fungi. To determine the scale at which the maps are most correlated (broad or fine-scale recombination rates), we further investigated how the correlations vary when various window sizes are used. We find that the correlations, consistently inferred with different correlation measures, peak at the 0.5-1 Mb scale (Fig. 7B), suggesting that the recombination landscape is conserved at large scales but shows rapid evolution at smaller scales. These results mirror findings from other eukaryotic species (e.g., [15,16]) and suggest that distinct mechanisms determine the recombination landscape at fine and broad scales in these two species.

**Figure 6:**
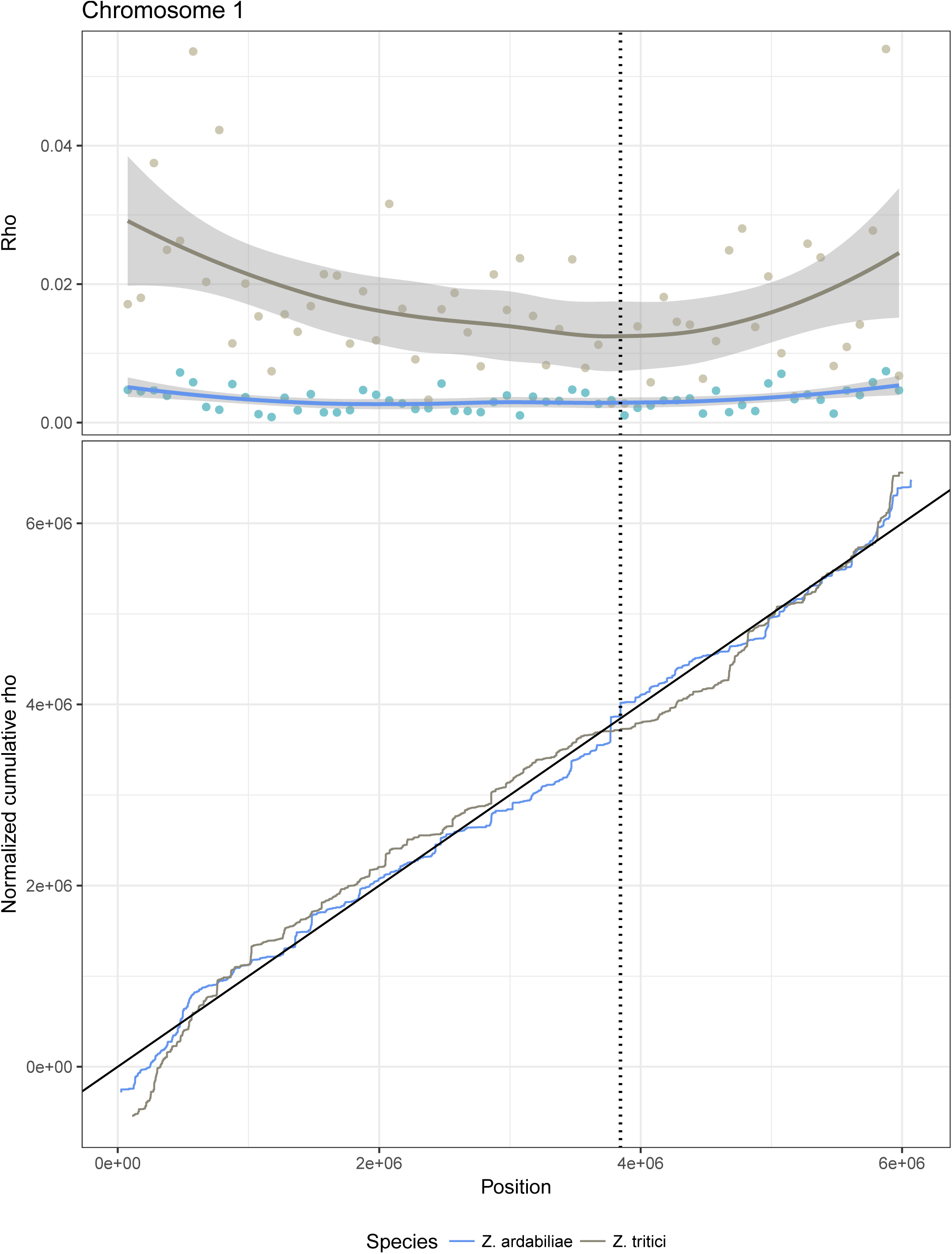
Recombination maps of *Z. tritici* and *Z. ardabiliae* plotted along the chromosome 1 of *Z. tritici*. A) Recombination map in 100 kb windows plotted together with smoothing curves. B) Cumulative curves of the recombination maps, scaled in order to be comparable. Figures for other chromosomes are available as Supplementary Data.

**Figure 7:**
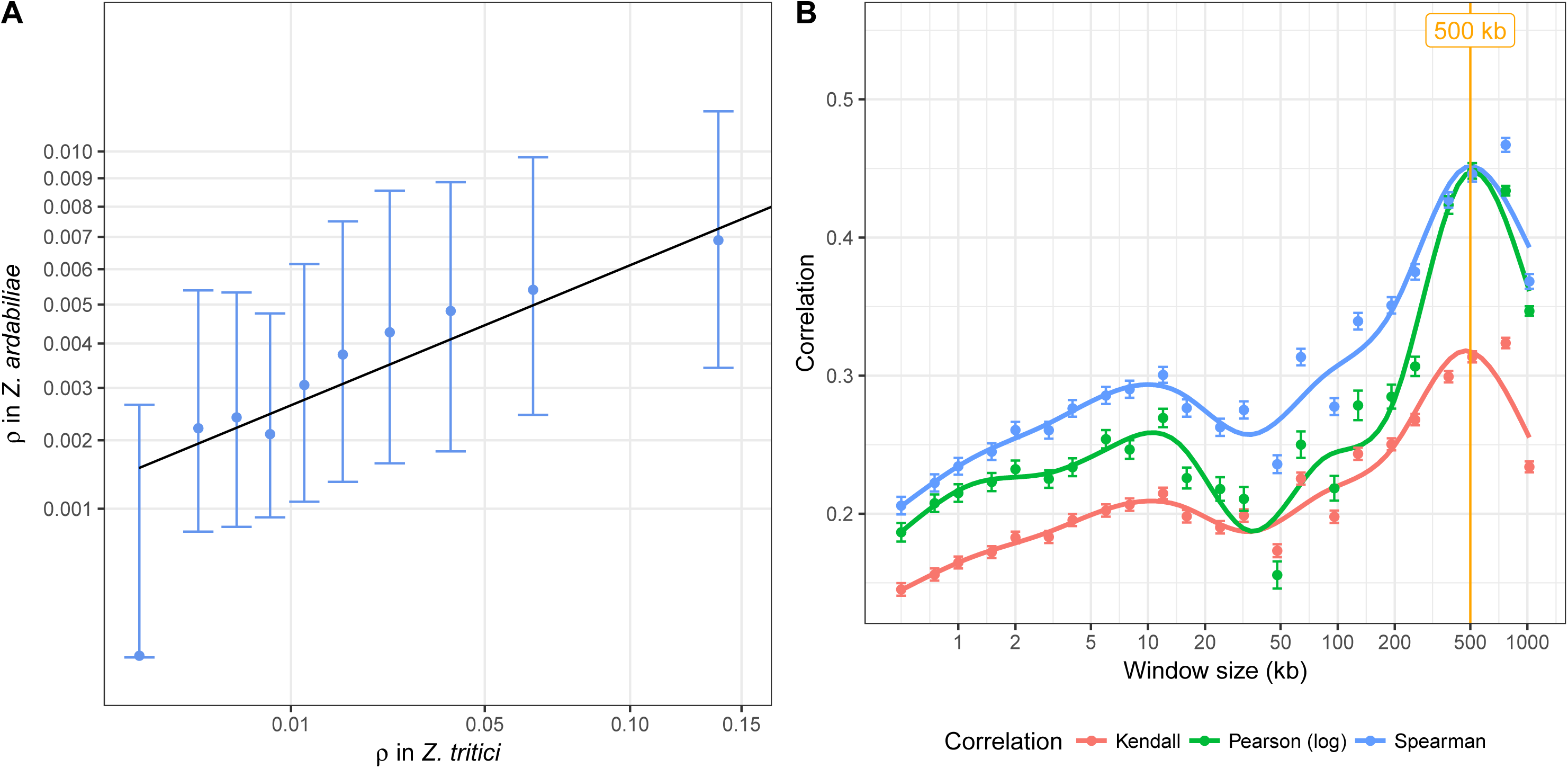
Correlation of recombination maps of *Z. tritici* and *Z. ardabiliae.* A) Comparison of the two recombination maps based on average recombination rates in windows of at least 100 SNPs in each species. Points represent averages in 10 classes with equal numbers of windows, points and error bars represent the median, first and third quartile of the distribution for each category. B) Correlation of recombination maps in sliding windows of different sizes. Three distinct correlation coefficients are plotted against recombination rates averaged in different window sizes (see Materials and Methods). Points indicate the averages of 1,000 samples and bars shows the standard errors of the means. Lines correspond to local regression smoothing (LOES).

### Frequency and intensity of recombination hotspots is higher in *Z. tritici*

The fine scale Ldhat recombination maps clearly reveal the presence of distinct peaks of recombination in both *Z. tritici* and *Z. ardabiliae* (Fig. 3). We used the program Ldhot to call positions of statistically significant recombination hotspots [76] and applied highly stringent selection criteria (see Materials and Methods) to obtain positions of the most significant hotspots for which the within-hotspot rate was at least ten times higher than the flanking regions (Fig. 8A). Interestingly, our approach revealed a considerably greater number of recombination hotspots in *Z. tritici* (2,578 hotspots) than in *Z. ardabiliae* (862 hotspots). Furthermore, we find a significant difference in the size of the hotspot regions between the two species. In general, the recombination hotspots span significantly shorter regions in *Z. tritici* (median 39 base pairs) than in *Z. ardabiliae* (66 base pairs, Wilcoxon ranked test p-value < 2.2e-16). We also compared the intensity of the recombination hotspots, as estimated by Ldhot (ρ across hotspot) and also find the median value of ρ in hotspots to be significantly higher in *Z. tritici* (median of 16.44 compared with 8.42 for *Z. ardabiliae*, Wilcoxon rank test p-value < 2.2e-6). The higher frequency of more intense hotspots in *Z. tritici* not only reveals a different hotspot landscape in the wheat pathogen; it also suggests that the overall higher recombination rate we observe in *Z. tritici* partly is explained by the different recombination hotspots architecture. While the differences to some extent can mirror the larger density of SNPs in *Z. tritici* that enables a finer resolution of the hotspot distribution and structure, we also speculate that recombination hotspots in these fungi have evolved since the divergence of *Z. tritici* and *Z. ardabiliae*. To address the extent of conservation in hotspot positions, we correlated the hotspot maps of the two species.

**Figure 8:**
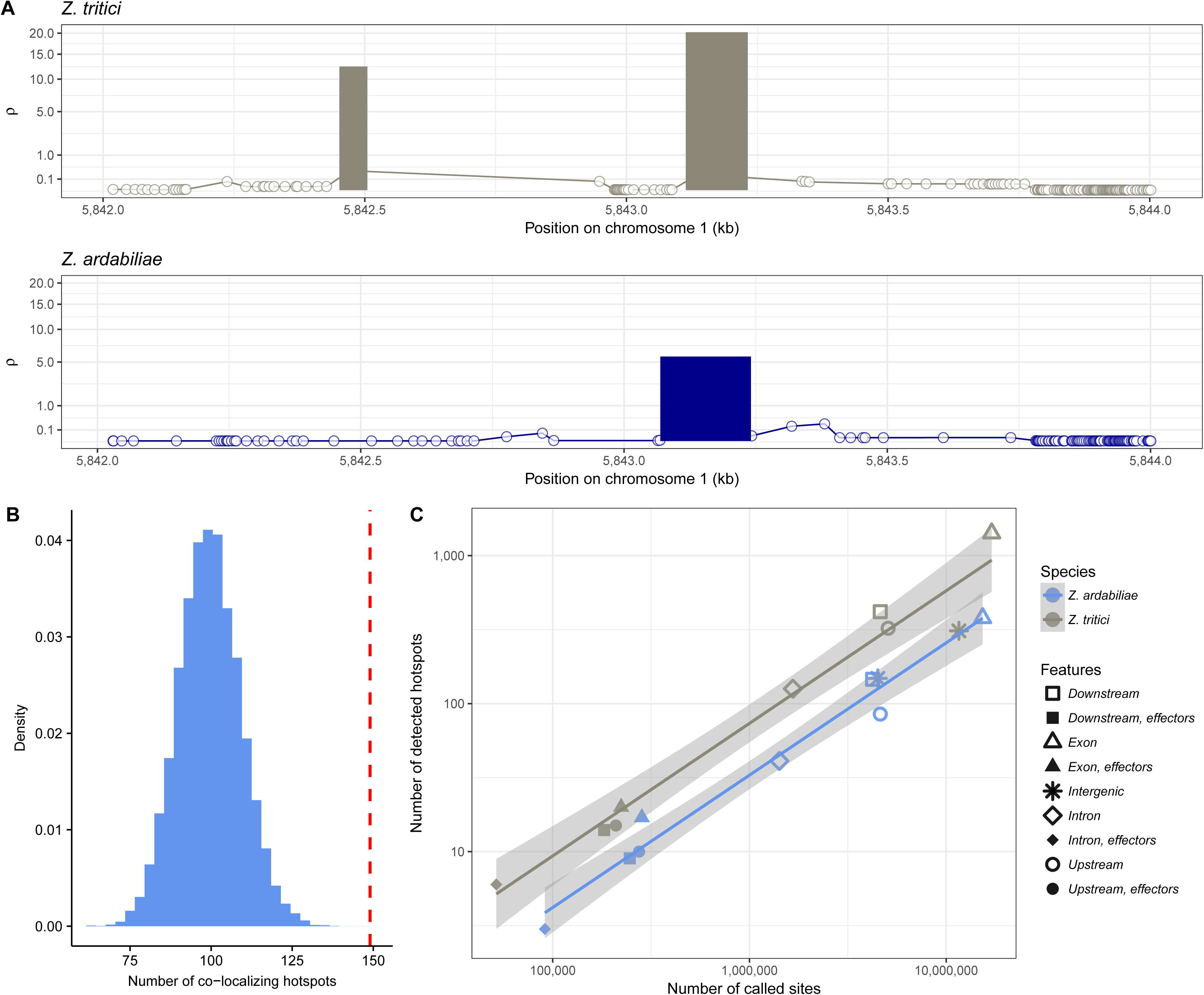
Distribution of hotspots in the genomes of *Z. tritici* and *Z. ardabiliae*. A: example mapped hotspot in a homologous region in *Z. tritici* and *Z. ardabiliae*. Lines indicate the background recombination rate as estimated by Ldhat. Bars indicate the positions, widths and strengths of hotspots detected by Ldhot in the region, after filtering (see Materials and Methods). B: Number of hotspots in *Z. tritici* in the direct 1 kb range of a hotspot in *Z. ardabiliae* (vertical line) and the corresponding distribution under the null hypothesis of a random distribution of hotspots. C: Frequencies of hotspots in distinct regions of the genome. Number of detected hotspots in each region as a function of the number of called sites. Lines correspond to ordinary least square regressions.

The position of recombination hotspots is defined by different mechanisms in different taxa, e.g. PRDM9 in primates and transcription start and end sites in other species such as birds [12,16]. Consequently, hotspot positions are highly conserved in some species [16,77], while highly variable in others [22]. We mapped *Z. ardabiliae* hotspots on the *Z. tritici* genomes and counted the numbers of co-localizing hotspots in the two species. We considered a hotspot in *Z. tritici* as co-localizing with a hotspot in *Z. ardabiliae* if the distance between the two hotspots is less than 1kb and if no other hotspot is present in between. We report that only 149 hotspots are co-localizing (6% of hotspots in *Z. tritici* and 20% of hotspots in *Z. ardabiliae*). This number is however significantly more than expected by chance (p-value < 9.99e-5, permutation test, Fig. 8B). These results are consistent with the previously reported genetic maps of *Z. tritici,* which also show little overlap of hotspots positions between two Swiss crosses [40]. Conversely, the patterns are highly different from *Saccharomyces* species in which hotspot positions are highly conserved and associated with functional elements across the yeast genomes [77].

Given the dense genomes of *Z. tritici* and *Z. ardabiliae* we assessed the number of hotspots mapped to coding sequences. Of the 2,578 *Z. tritici* hotspots, 132 are located in introns and 1,435 are located in exons. Interestingly, in *Z. ardabiliae* we find 44 hotspots in introns and only 396 in exons. We plotted the number of hotspots as a function of the number of called sites in each region (Fig. 8C). We observe a general trend in which the number of detected hotspots increases with the number of called sites as a power law (linear relationship in log space), and with more hotspots detected in *Z. tritici*. In contrast to patterns of previously studied species, this reveals the presence of hotspots in all parts of the genome, including coding regions. We do not observe a significant enrichment close to transcription start site (upstream regions) like in yeast [23]. We further note that comparatively fewer hotspots locate in intergenic regions of *Z. tritici*, these regions displaying a density of hotspots similar to what is expected in *Z. ardabiliae* for the observed number of callable sites. We hypothesize two non-exclusive possible origins for this result: (1) the number of callable sites is higher in *Z. tritici* intergenic regions than in *Z. ardabiliae*, due to the lack of telomere-to-telomere assembly of a reference genome for this species. The missing regions could potentially bias our estimate of hotspot densities in intergenic regions. (2) another possible explanation is that the comparatively larger number of hotspots in *Z. tritici* is due to an increased hotspot density in protein-coding genes in this species, which raises the question whether intragenic recombination hotspots represent a selected feature during evolution of the wheat-infecting lineage.

## Conclusions

Pathogens need to adapt rapidly to overcome immune responses in their host [78]. Several examples from animal and plant pathogens document exceptionally high rates of genome rearrangements including changes in ploidy and full chromosome gains or losses (e.g., [79–82]). So far the importance of meiotic recombination in rapid evolution of pathogens has been poorly addressed. Our analyses demonstrate extraordinary high recombination rates in two fungal plant pathogens and thereby suggest that sexual recombination also can be a major driver of rapid pathogen evolution.

The overall higher recombination rate and the increased density of recombination hotspots in the crop pathogen *Z. tritici* are remarkable. *Z. tritici* and *Z. ardabiliae* share a recent common ancestor, but exist and evolve in highly different environments. While *Z. ardabiliae* infects wild grasses in a natural ecosystem, *Z. tritici* infects a crop host and propagate only in managed ecosystems. Agricultural management strategies, dense host populations and increased gene flow between geographically distant populations are factors that contribute to a different population structure of *Z. tritici*. We hypothesize that an increased rate of recombination in coding sequences of *Z. tritici* was selected as it favored the rapid generation of new alleles and allele combinations [83]. The exceptionally high recombination rate in *Z. tritici* allows the pathogen to rapidly overcome new host resistances and explains the current difficulties of controlling this important wheat pathogen.

## Materials and methods

### Genome data

The lifecycle of *Z. tritici* is predominantly haploid and the genome analyses conducted here thus rely on haploid genome data. The 40-Mb reference genome of the *Z. tritici* isolate IPO323 was sequenced at the Joint Genome Institute using Sanger sequencing [38]. Two Iranian *Z. tritici* isolates and four Iranian *Z. ardabiliae* isolates were sequenced in a previous study using Illumina sequencing (Table S1) [37]. We used genome data from an additional ten isolates of *Z. tritici* that originate from wheat fields in Denmark, France and Germany [44]. In this study, we report the genome sequences of thirteen isolates of *Z. ardabiliae* that originate from wild grasses collected in the province of Ardabil in Iran (Table S1). DNA extraction was performed as previously described [37]. Library preparation and paired end sequencing using an Illumina HiSeq2000 platform were conducted at Aros, Skejby, Denmark. Sequence data has been deposited under the NCBI BioProject IDs PRJNA277174.

The thirteen re-sequenced *Z. ardabiliae* genomes were assembled from 100 bp paired end reads using the de novo assembly algorithm of the CLC Genomics Workbench version 5.5 (Qiagen, Aarhus, Denmark). The assemblies were created using standard settings for paired-end reads. We used a previously published RNAseq based annotation to distinguish the parameter estimates for coding and non-coding sequences [39]. To predict the genes that encode effectors we used the software EffectorP [75] with default settings, on genes predicted by SignalP [84] to encode a secreted protein.

### Genome alignment and SNP calling

Genome alignments were separately created for each population using the MultiZ program from the TBA package [85]. Default parameters were used, although LastZ was used instead of BlastZ for pairwise alignments. Genome alignments were projected against the two reference genomes of each species: IPO123 for *Z. tritici* and STO4IR-1.1.1 for *Z. ardabiliae [37,38]*. The projected alignments in MAF format were filtered using the MafFilter program [86] with the following filters: 1) each syntenic block was realigned using Mafft [87], and blocks with more than 10 kb were split for computational efficiency; 2) only blocks where all individuals were present were retained (13 *Z. tritici* and 17 *Z. ardabiliae*); 3) a window of 10 bp was slid by 1 bp, and windows containing at least one position with gaps in at least 2 species were discarded and the containing blocks were split; 4) a window of 10 bp was slid by 1 bp, and windows with a total of more than 100 gaps were discarded and the containing blocks were split; and 5) all blocks were merged according to the reference genome with empty positions filled by 'N', which resulted in one masked alignment per chromosome for *Z. tritici* and one masked alignment per contig for *Z. ardabiliae*. The chromosome and contig alignments were further divided in non-overlapping windows of 1 Mb (data set 1) or 100 kb (data set 2). The MafFilter program was further used to estimate statistics on the alignments at each filtering step, and to compute the nucleotide diversity (Watterson's θ) from the final filtered genome alignments.

### Estimating recombination

Filtered alignments (1-Mb windows, data set 1) were exported as fasta files for the Ldhat and Ldhelmet packages. The program *convert* from the Ldhat package was used to convert fasta files into input loci files for the program *interval [41]*. Only fully resolved biallelic positions were exported (see Table 1 for the details of SNP numbers). Likelihood tables were generated for θ values of 0.0005, 0.005 and 0.05. The *interval* program was run with 10,000,000 iterations and sampled every 5,000 iterations with a burn-in of 100,000 iterations. Ldhelmet was run with the parameters suggested in the user manual [13] and <u>https://sourceforge.net/projects/ldhelmet/</u>). Comparison of recombination maps on the same set of SNPs was performed using standard principal component analysis, as implemented in the R package ade4 [88]. A table was computed with one column per method (LDhat and LDhelmet, each with theta set to 0.0005, 0.005 or 0.05) and one line per analyzed SNP pair, and the two first principal components kept to plot a correlation circle (Figure 1A).

In order to assess the robustness of the recombination maps, alternative maps for *Z. tritici* were constructed using the same protocol (1) after discarding all singletons, (2) after removing five individuals to ensure absence of population structure. All maps were compared to the previously published genetic map of Croll et al [40] in windows of 20 kb. Correlations were assessed using Kendall´s rank correlation test, and confidence intervals were obtained by bootstrapping windows.

We calculated average recombination rates in windows and regions by taking the average of recombination estimates between every pairs of SNPs, weighted by the physical distance between the SNPs. Pairs of SNPs for which the confidence interval of the recombination estimate was higher than two times the mean were discarded and therefore not used in the average computation. Using the gene annotations available for the two reference species [39], we calculated the following information for each gene: 1) the average recombination rate in exons, 2) the average recombination rate in introns, and 3) the average recombination rate in the 500 bp flanking 5' region and 4) in the 500 bp flanking 3' region. We also calculated the average recombination rate for each intergenic region (500 bp from / to genes). GFF3 files from [39] were retrieved and processed using the “Genome Tools” package to add intron annotations [89]. The resulting gene annotations were analyzed in R together with recombination maps [90].

### Assessment of LD-based recombination estimates by simulation

We used the SCRM coalescent simulator [91] in order to simulate polymorphism data with a constant mutation rate but variable recombination rate. Recombination rates were drawn randomly from an exponential distribution with mean 0.02. Segments with piecewise constant recombination rate were taken randomly from an exponential distribution with mean 100 kb. Sample sizes of 10, 30 and 100 individuals were tested for comparison, with a population mutation rate equal to 0.05, 0.005, 0.0005 and 0.00005. We generated a locus of 10 Mb for simulations with θ equal to 0.005, 0.0005 and 0.00005, but only 1 Mb for simulations with θ equal to 0.05, as the resulting output file from LDhat would otherwise become excessively large due to the high number of SNPs. The true recombination rate used at each position of the alignment was recorded for later comparison. The output of SCRM was converted to Ldhat input format using python scripts. Recombination rates were estimated using the interval program from the Ldhat package [41]. For simulations with θ = 0.05 and 0.005 a likelihood lookup table with θ = 0.01 was used, whereas a lookup table with θ = 0.001 was used for simulations with θ = 0.0005 and 0.00005. The inferred recombination rate at each position was then compared to the true rate. A variant of this simulation procedure was used to assess the impact of population structure on the inference of recombination rate. The SCRM coalescent simulator was used with a five-islands population model, with sample sizes 2, 3, 4, 5 and 6 per deme, resulting in a total of 20 individuals. Migration rates were assumed to be all identical between demes, and values of M = 4Ne.m = 1, 10 and 100 were tested. 1Mb regions were simulated with θ = 0.005 for each migration rate.

### Reference species alignment and comparison

The two reference strains IPO323 (*Z. tritici*) and ST11IR-11.4.1 (*Z. ardabiliae*) were aligned using LastZ [85]. The resulting genome alignment was used to map the coordinates of *Z. ardabiliae* SNPs to the *Z. tritici* genome, using the MafFilters “lift-over” filter [86]. A total of 893,171 (86%) positions could be mapped from *Z. ardabiliae* to *Z. tritici* and were used for further analyses. Non -overlapping windows containing at least 100 analysed SNPs in each species were generated for the comparison of recombination rates between the two species.

### Multi-scale correlations

We calculated the average recombination rates in windows of varying sizes and retained only windows that contained at least 1% of the polymorphic positions. To enforce a similar statistical power among different window sizes, a number of windows were chosen randomly. The same number of randomly chosen windows was used for the distinct comparisons. To assess the sampling variance, 1,000 independent samplings (with replacement) were performed for each window size. Window sizes of 0.5, 1, 2, 4, 8, 16, 32, 64, 128, 256, 512 and 1,024 kb were tested, with 27 windows sampled in each case. We measured correlation coefficients using the Spearman, Kendall and Pearson’s correlation coefficients. Spearman and Kendall’s coefficients are ranked-based; therefore they do not assume bi-normality as Pearson’s coefficient does. Because recombination rates are typically exponentially distributed, Pearson’s coefficient was measured for the log rates instead of the raw ρ rates. Spearman’s coefficient assumes that the variables are continuously distributed; therefore it does not resolve ties. Thus jittering was used to randomly resolve ties in the input variables (R function ‘jitter’, with default parameters). Conversely, Kendall’s coefficient assumes ordinal input variables. Therefore, using the three correlation measures allows assessing the robustness of the correlation signal. A graphical representation was performed using the ggplot2 package for R, which performed local polynomial regression fitting for the curves [92].

### Mapping of hotspots

Hotspots were detected using the Ldhot program [76]. For computational efficiency, Ldhot was run on the 100 kb alignments (data set 2). A background recombination map was first estimated for each alignment using the *interval* program of Ldhat with a θ value of 0.005 [41]. The resulting maps were highly correlated with the maps based on 1-Mb alignments and showed little effect of the discretization scheme. The background recombination map was used as input to Ldhot with default parameter values and 1,000 simulations.

Significant hotspots were filtered for further analysis. First, only the hotspots with a value of ρ between 5 and 100 across the hotspot coordinates were selected because higher values are most likely artifacts and the performance of Ldhot is low for weak hotspots [76]. A few hotspots with extremely large sizes (> 2 kb) were further discarded. This process identified 9,133 hotspots in *Z. tritici* and 1,287 hotspots in *Z. ardabiliae*. We calculated the mean background rate in each detected hotspot and in the two 20-kb flanking regions. We further selected hotspots for which the within-hotspot rate was at least ten times higher than the flanking regions. Thus 2,578 and 862 hotspots were identified in *Z. tritici* and *Z. ardabiliae,* respectively. The *Z. ardabiliae* hotspots were mapped onto the *Z. tritici* genome using MafFilter’s liftover function [86]. We considered a hotspot in *Z. tritici* as co-localizing with a hotspot in *Z. Ardabiliae* if the distance between them was less than 1kb, and if no other hotspot was found between the two. We compared statistics on the distribution of hotspots by randomizing the hotspot positions while keeping their original size, for each chromosome independently. In order to do so, we used the following procedure:

1) compute the total “inter-hostpots” distance, L, as the sum of all distances between consecutive hotspots,

2) draw random distinct positions uniformly in [1, L]. These positions are the starting positions of each randomized interval,

3) order, then expand each interval to match its original size and compute the corresponding end positions. Correct the coordinates in order to account for previous intervals.

In order to account for variable coverage along the genome, we also simulated intervals corresponding to chromosome regions that were not included in our analysis, using the same procedure as for hotspots randomization. Each randomized set of hotspots therefore contains the same amount of callable sites as the actual analysis. We assessed the significance of the number of co-localizing hotspots using 10,000 permutations. The corresponding R scripts are available as Supplementary Data 3.

### Models of GC content evolution

The two reference strains IPO323 (*Z. tritici*) and ST11IR-11.4.1 (*Z. ardabiliae*) were aligned using LastZ [85]. Several filtering steps were further applied to the alignment. First, each synteny block was realigned using the MAFFT aligner [87] after splitting block longer than 10 kb for computational efficiency, which resulted in an alignment of 27,918,318 bp that included both species. Second, a window of 30 bp was slid by 1 bp along the alignment. Windows with more than 29 gaps in total between the two species were further discarded, which resulted in 27,237,601 filtered positions. To minimize the effect of selection on GC patterns, we further discarded regions in the alignment that were annotated as protein-coding genes in one or both species. This resulted in a total alignment of 9,143,114 bp. The alignment was further divided into windows ranging from 1 to 4 kb and only data from the essential chromosomes (*Z. tritici* chromosomes 1 to 13) were retained. The final alignment contained 2,052 cleaned windows containing sequences for both species with no synteny break, and it encompassed 3,179,581 bp. A model of sequence evolution was independently fitted on each window using maximum likelihood [93]. The HKY85 model was used as a basis allowing three frequency parameters ((G + C) / (A + C + G +T), A / (A + T) and G / (G + C)) in addition to the transition over transversion ratio [94]. We fitted a non-homogeneous, non-stationary model of substitution, allowing us to estimate three distinct GC contents for *Z. tritici*, *Z. ardabiliae* and their common ancestor. Other parameters were considered constant between species and their ancestor. A molecular clock was assumed (so that the two branches leading to *Z. tritici* and *Z. ardabiliae* were equal in length) and a 4 classes gamma distribution of rates with a shape parameter fixed to 0.5 was used. We further calculated the observed GC content in each species for each window. The average recombination rate was calculated for each windows containing at least 1% polymorphic position (leaving 1,642 windows).

As similar analysis was conducted using recombination rate estimated from [40] which were calculated in 20 kb windows. The corresponding pairwise alignment regions were extracted and filtered, and coding regions from both species were discarded, which resulted in 1,948 windows of at least 1 kb where a non-homogeneous, non-stationary model of substitution could be fitted.

## Availability of Data and Materials

Illumina reads for *Z. ardabiliae* are available from NCBI under the Biosample IDs SAMN05818736-SAMN05818752.

Illumina reads for *Z. tritici* are available from NCBI under the BioProject ID PRJNA312067. FigShare repository for scripts 10.6084/m9.figshare.3806244.

## Acknowledgements

EHS is supported by intramural funding from the Max Planck Society, Germany, a personal grant from the State of Schleswig-Holstein, Germany and a grant from the German Research Council, DFG, grant number HO 4435/1-1. JYD is supported by intramural funding from the Max Planck Society, Germany. The authors thank Nicolas Galtier for helpful discussions on the gBGC and Daniel Croll for providing genetic data from experimental crosses of *Z. tritici*.

## Supplementary Material

**Table S1:** Summary information of *Z. tritici* and *Z. ardabiliae* isolates used in the study.

**Figure S1:**
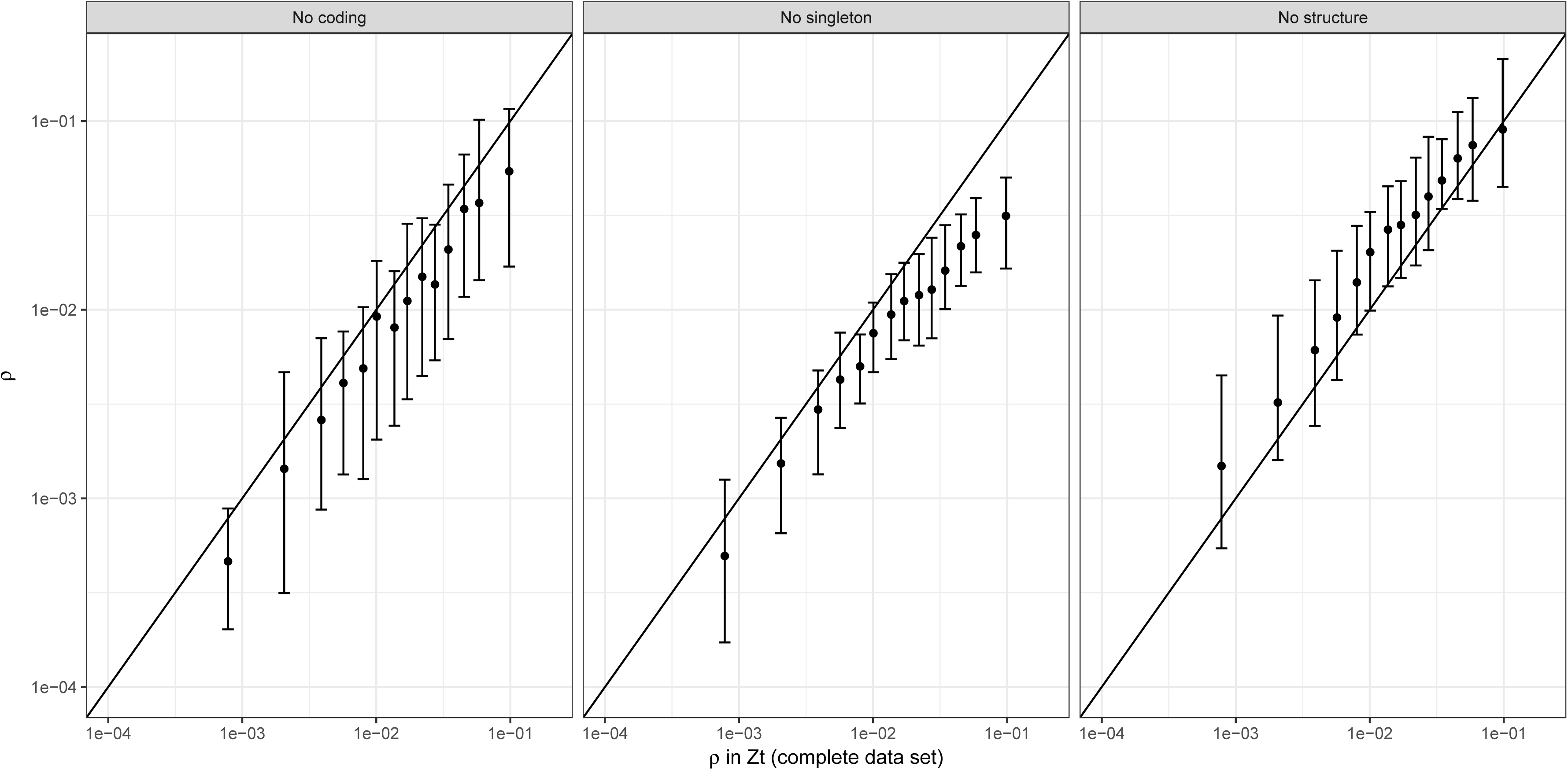
Effect of selection, sequencing errors and population structure on the inference of population recombination rate. Three population recombination maps (y-axis) are compared to the map used throughout this study (x-axis). Left panel recombination map obtained after removing SNPs within and nearby genes. Middle panel: recombination map obtained after discarding all singletons. Right panel: recombination map obtained after keeping only eight strains showing no signature of population structure.

**Figure S2:**
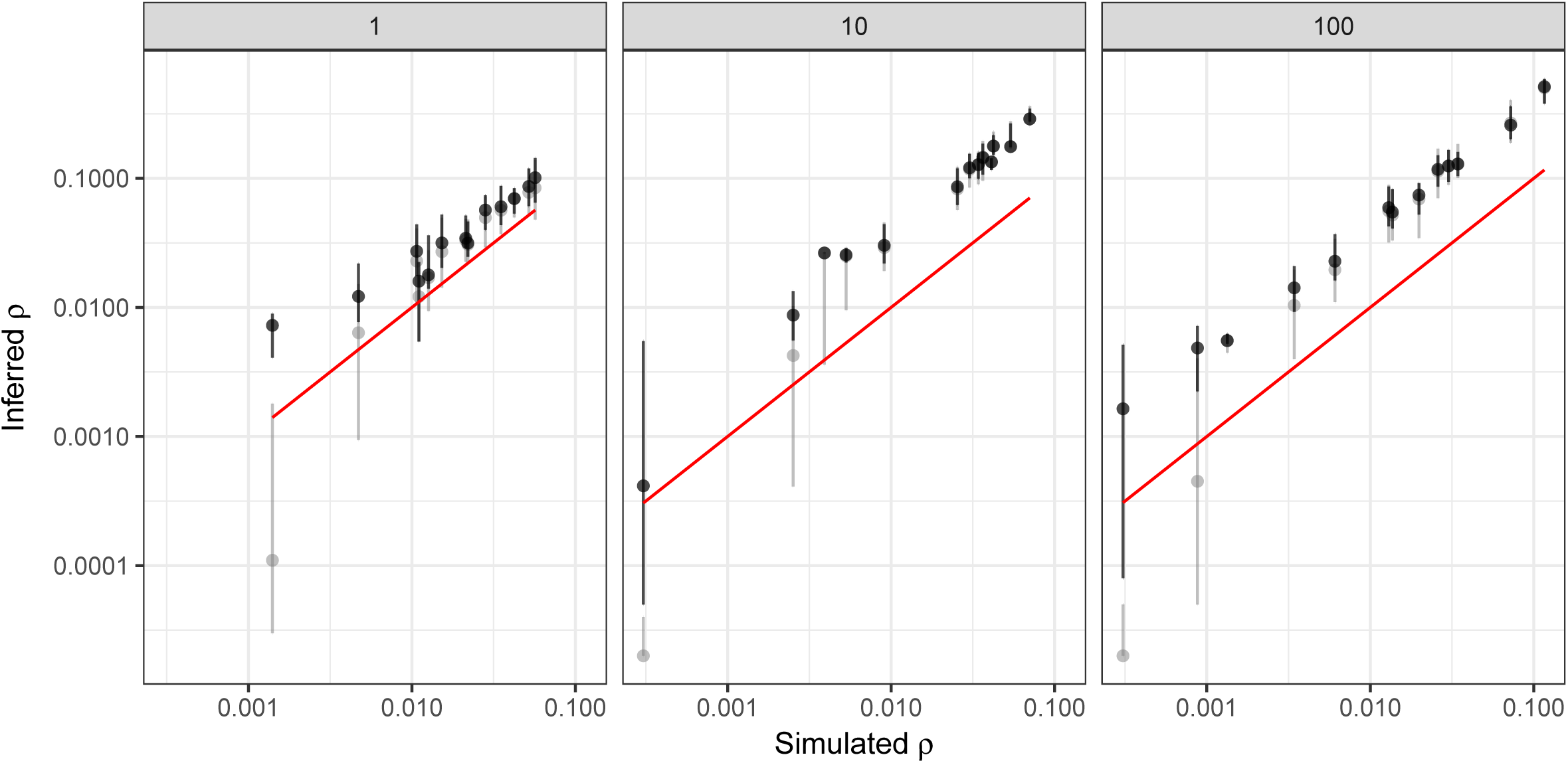
Effect of population structure on the inference of population recombination rate. 1 Mb regions were simulated as in Figure 2, with θ = 0.005 and n = 20 individuals. A five islands model of population structure was used, with sample sizes 2, 3, 4, 5 and 6 and a scaled migration rate M = 4Ne.m as indicated in each panel. Legends as in Figure 2.

**Figure S3:**
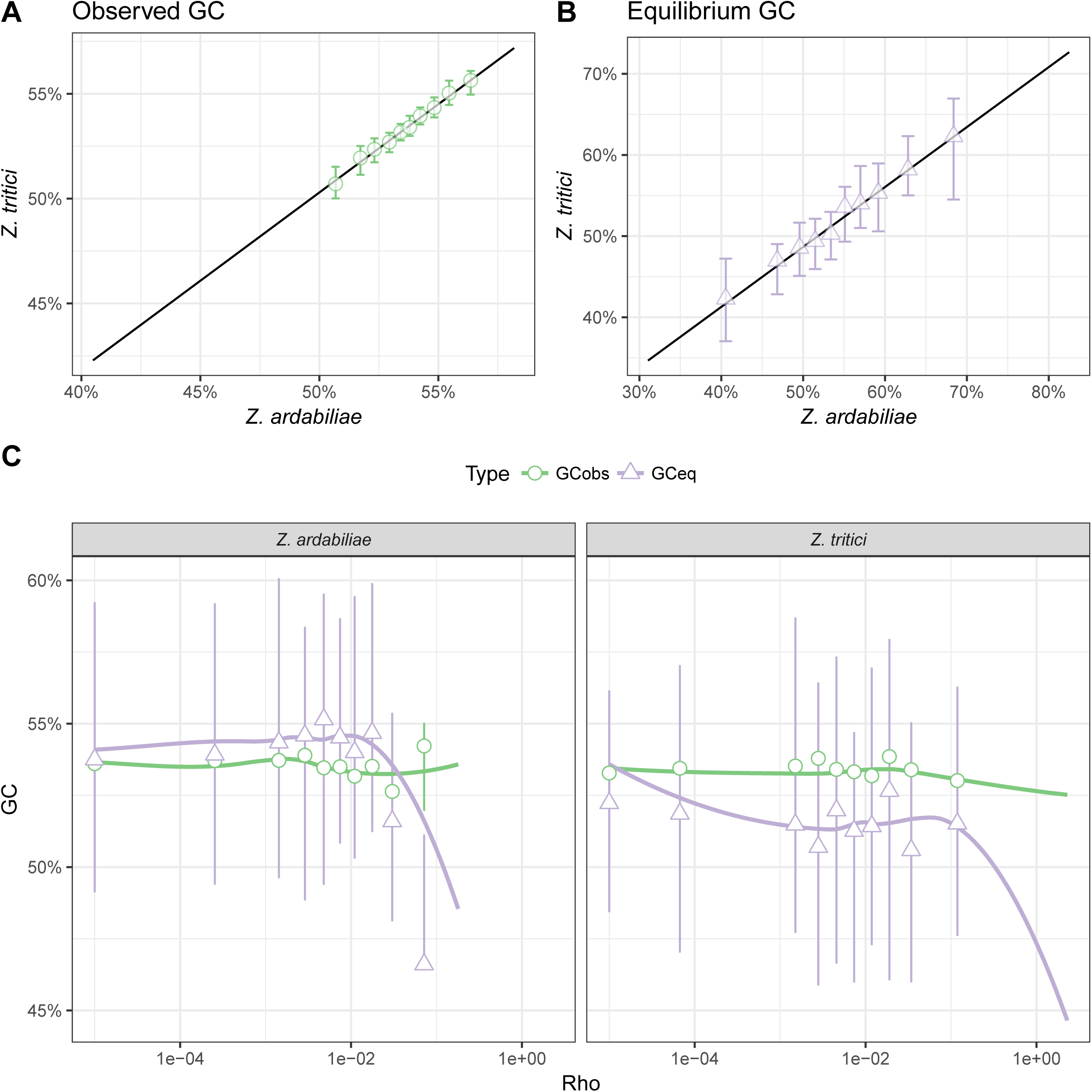
Genome-wide recombination rate and GC content. A) Observed GC content in *Z. tritici* plotted against observed GC content in *Z. ardabiliae*. B) Equilibrium GC content in *Z. tritici* plotted against equilibrium GC content in *Z. ardabiliae*. C) GC content as a function of recombination rate. For all three panels, the x-axis was discretized in 10 categories with the same amount of data. Points indicate the median GC content in each category; and bars correspond to the first and third quartiles of the distribution. For panels A and B, the 50% quantile regression line is shown as a straight line. For panel C, curves represent local regression smoothing (LOES). In the three cases, the lines and curves where fitted to the data without discretization.

**Supplementary Data 1:** Chromosomal patterns for every chromosomes. Legends as for Figure 3.

**Supplementary Data 2:** Correlation of recombination maps for every chromosome. Legends as for Figure 6.

**Supplementary Data 3:** All scripts and data allowing reproducing results and figures in this manuscript.

